# ElectroPhysiomeGAN: Generation of Biophysical Neuron Model Parameters from Recorded Electrophysiological Responses

**DOI:** 10.1101/2023.12.19.572452

**Authors:** Jimin Kim, Minxian Peng, Shuqi Chen, Qiang Liu, Eli Shlizerman

## Abstract

Recent advances in connectomics, biophysics, and neuronal electrophysiology warrant modeling of neurons with further details in both network interaction and cellular dynamics. Such models may be referred to as ElectroPhysiome, as they incorporate the connectome and individual neuron electrophysiology to simulate neuronal activities. The nervous system of *C. elegans* is considered a viable framework for such ElectroPhysiome studies due to advances in connectomics of its somatic nervous system and electrophysiological recordings of neuron responses. In order to achieve a simulated ElectroPhysiome, the set of parameters involved in modeling individual neurons need to be estimated from electrophysiological recordings. Here, we address this challenge by developing a deep generative estimation method called ElectroPhysiomeGAN (EP-GAN), which once trained, can instantly generate parameters associated with the Hodgkin-Huxley neuron model (HH-model) for multiple neurons with graded potential response. The method combines Generative Adversarial Network (GAN) architecture with Recurrent Neural Network (RNN) Encoder and can generate an extensive number of parameters (>170) given the neuron’s membrane potential responses and steady-state current profiles. We validate our method by estimating HH-model parameters for 200 simulated neurons with graded membrane potential followed by 9 experimentally recorded neurons (where 6 of them newly recorded) in the nervous system of *C. elegans*. Comparison of EP-GAN with existing estimation methods shows EP-GAN advantage in the accuracy of estimated parameters and inference speed for both small and large number of parameters being inferred. In addition, the architecture of EP-GAN permits input with arbitrary clamping protocols, allowing inference of parameters even when partial membrane potential and steady-state currents profile are given as inputs. EP-GAN is designed to leverage the generative capability of GAN to align with the dynamical structure of HH-model, and thus able to achieve such performance.

## Introduction

Models of the nervous system aim to achieve biologically detailed simulations of large-scale neuronal activity through the incorporation of both structural connectomes (connectivity maps) and individual neural dynamics. The nervous system of *Caenorhabditis elegans* (*C. elegans*) is considered a framework for such a model as the connectome of its somatic nervous system for multiple types of interaction is mapped [1, 2, 3]. In addition to the connectome, advances in electrophysiological methodology allow the recording of whole-cell responses of individual neurons. These advances provide biophysically relevant details of individual neuro-dynamical properties and warrant a type of model for the *C. elegans* nervous system incorporating both the connectomes and individual biophysical processes of neurons. Such a model could be referred to as *ElectroPhysiome*, as it incorporates a layer of individual neural dynamics on top of the layer of inter-cellular interactions facilitated by the connectome.

The development of nervous system models that are further biophysically descriptive for each neuron, i.e., modeling neurons using the Hodgkin-Huxley type equations (HH-model), requires fitting a large number of parameters associated with ion channels found in the system. For a typical single neuron, these parameters could be tuned via local optimizations of individual ion channel parameters estimated separately to fit their respective *in-vivo* channel recordings such as activation/inactivation curves [4, 5, 6, 7, 8, 9]. Such method requires multiple experiments to collect each channel data and when such experiments are infeasible, the parameters are often estimated through hand-tuning. In the context of developing the ElectroPhysiome of *C. elegans*, the method would have to model approximately 300 neurons each including an order of hundreds of parameters associated with up to 15 to 20 ionic current terms (with some of them having unknown ion channel composition), which would require large experimental studies [7]. Furthermore, the fitted model may not be the unique solution as different HH-parameters can produce similar neuron activity [10, 11, 12, 13]. As these limitations also apply for general neuron modeling tasks beyond *C. elegans* neurons, there has been an increasing search for alternative fitting methods requiring less experimental data and manual interventions.

A promising direction in associating model parameters with neurons has been the simultaneous estimation of all parameters of an individual neuron given only electrophysiological responses of cells, such as membrane potential responses and steady-state current profiles. Such an approach requires significantly less experimental data per neuron and offers more flexibility with respect to trainable parameters. The primary aim of this approach is to model macroscopic cell behaviors in an automated fashion. Indeed, several methods adopting the approach have been introduced. Buhry et al 2012 and Laredo et al 2022 utilized the Differential Evolution (DE) method to simultaneously estimate the parameters of a 3-channel HH-model given a whole-cell membrane potential responses recording [14, 15]. Naudin et al 2022 further developed the DE approach and introduced the Multi-objective Differential Evolution (DEMO) method to estimate 22 HH-parameters of 3 non-spiking neurons in *C. elegans* given their whole-cell membrane potential responses and steady-state current profiles [16]. The study was a significant step toward modeling whole-cell behaviors of *C. elegans* neurons in a systematic manner. From a statistical standpoint, Wang et al 2022 used the Markov-Chain-Monte-Carlo method to obtain the posterior distribution of channel parameters for HH-models featuring 3 and 8 ion channels (2 and 9 parameters respectively) given the simulated membrane potential responses data [17]. From an analytic standpoint, Valle et al 2022 suggested an iterative gradient descent based method that directly manipulates HH-model to infer 3 conductance parameters and 3 exponents of activation functions given the measurements of membrane potential responses [18]. Recent advances in machine learning gave rise to deep learning based methods which infer steady-state activation functions and posterior distributions of 3-channel HH-model parameters inferred by an artificial neural network model given the membrane potential responses data [19, 20].

While these methods suggest that simultaneous parameter estimation from macroscopic cell data is indeed possible through a variety of techniques, it is largely unclear whether they can be extended to fit more detailed HH-models featuring a large number of channels and parameters (e.g., *C. elegans* neurons) [7]. Furthermore, for most of the above methods, the algorithms require an independent (from scratch) optimization process for fitting each individual neuron, making them difficult to scale up the task toward a large number of neurons.

Here we propose a new machine learning approach that aims to address these aspects for the class of non-spiking neurons, which constitute the majority of neurons in *C. elegans* nervous system [21]. Specifically, we develop a deep generative neural network model (GAN) combined with a recurrent neural network (RNN) encoder called ElectroPhysiomeGAN (EP-GAN), which directly maps electrophysiological recordings of a neuron, e.g., membrane potential responses and steady-state current profiles, to HH-model parameters of arbitrary dimensions (Figure 1). EP-GAN can be trained with simulation data informed by a generic HH-model encompassing a large set of arbitrary ionic current terms and thus can generalize its modeling capability to multiple neurons. Unlike typical GAN architecture trained solely with adversarial losses, we propose to implement additional regression loss for reconstructing the given membrane potential responses and current profiles from generated parameters, thus improving the accuracy of the generative model. In addition, due to the RNN component of EP-GAN, the approach supports input data with missing features such as incomplete membrane potential responses and current profiles.

**Figure 1:**
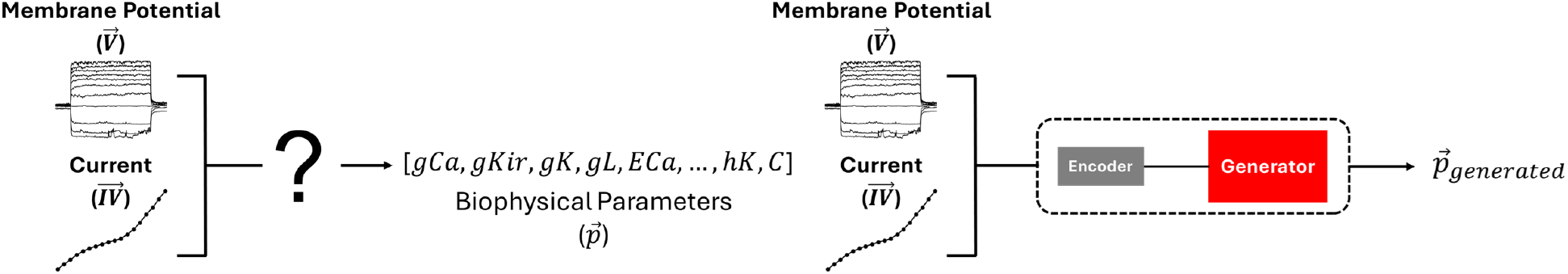
Estimation of HH-model parameters from membrane potential and steady-state current profiles. Given the membrane potential responses (V) and steady-state current profiles (IV) of a neuron, the task is to predict biophysical parameters of Hodgkin-Huxley type neuron model (Left). We use Encoder-Generator approach to predict the parameters (Right)

We validate our method to estimate HH-model parameters of 200 simulated non-spiking neurons followed by applying it to three previously recorded non-spiking neurons of *C. elegans*, namely RIM, AFD, and AIY. Studies have shown that membrane potential responses of these neurons can be well modeled with typical HH-model formulations with 22 parameters [16, 22]. We show that when trained with a more detailed HH-model consisting of 16 ionic current terms resulting in

175 trainable parameters, EP-GAN can predict parameters reproducing their membrane potential responses with higher accuracy in the reconstruction of membrane potential with significantly faster inference speed than existing algorithms such as Multi-Objective DE and Genetic Algorithms. Through ablation studies on input data, we show that EP-GAN retains its prediction capability when provided with incomplete membrane potential responses and steady-state current profiles. We also perform ablation studies on the model architecture components to elucidate each component’s contributions toward the accuracy of the predicted parameters. To further test EP-GAN, we estimate HH-model parameters for 6 newly recorded non-spiking *C. elegans* neurons: AWB, AWC, URX, RIS, DVC, and HSN, whose membrane potential responses were not previously modeled.

Our results suggest that EP-GAN can learn a translation from electrophysiologically recorded responses and propose projections of them to parameter space. EP-GAN method is currently limited to non-spiking neurons in *C. elegans* as it was designed and trained with HH-model describing the ion channels of these neurons. EP-GAN applications can be potentially extended toward resolving neuron parameters in other organisms since non-spiking neurons are found within animals across different species [23, 24, 25, 26, 27, 28, 29, 30].

## Results

We evaluate EP-GAN with respect to 4 existing evolutionary algorithms introduced for general parameter estimation: NSGA2, DEMO, GDE3, and NSDE. Specifically, NSGA2 is a variant of the Genetic Algorithm that uses a non-dominated sorting survival strategy, and is a commonly used benchmark algorithm for multiobjective optimization problems that include HH model fitting [31, 32, 33]. DEMO, GDE3, and NSDE are variants of multi-objective differential evolution (DE) algorithms that combine DE mutation with pareto-based ranking and crowding distance sorting applied in NSGA2’s survival strategy [34, 35, 36]. These methods have been proposed as more effective methods than direct DE for estimation of HH-model parameters [16, 37, 38, 39]. In particular, DEMO has been successfully applied to estimate HH-model parameters for non-spiking neurons in *C. elegans* [16]. All 4 methods support multi-objective optimization over large parameter space allowing them to have similar setups as EP-GAN. All 4 methods were implemented in Python where DEMO uses the algorithm proposed in [16] whereas NSGA2, GDE3 and NSDE were implemented using Pymoo package [40].

For the HH-model to be estimated, we use the formulation introduced in [7, 41]. The model features 16 ion channels that were found in *C. elegans* and other organisms expressing homologous channels and is considered the most detailed neuron model for the organism. The model has a total of **203** parameters, of which we identify 175 of them have approximate ranges with lower and upper bounds that can be inferred from the literature [8, 22, 42]. We thus target these 175 parameters as trainable parameters for all methods. For a detailed list of all 203 parameters included in the model and 175 parameters used for training, see [41] and the included table *predicted parameters* in supplementary files.

### Predictions on simulated neurons

We first validate EP-GAN by training and testing using simulated neurons. Each simulated neuron training sample consists of two inputs: i.) Simulated membrane potential traces concatenated with associated external stimuli traces and ii.) Steady-state currents across 18 voltage points. For each neuron, the output is the set of 175 HH-parameters to be inferred. Each membrane potential trace is simulated for 15 seconds for a given stimulus according to current-clamp protocol where the stimulation is applied for 5 seconds at [5, 10] seconds and no stimulation is applied at *t* = [0, 5) and *t* = (10, 15]. These time intervals are chosen to ensure sufficient stabilization periods before and after stimulation. For the membrane potential input, the responses during *t* = (4, 11) interval are used consistent with the time interval used by experimental recordings. Similarly, steady-state currents are computed across 18 voltage states according to voltage-clamp protocol (see Table 4 for detailed current/voltage clamp protocols used for simulated neurons). The output HH-parameters are of the simulated neurons chosen randomly from lower and upper bounds as previously described. For training EP-GAN, we simulate a total of 32,000 (32k) neurons where EP-GAN achieves both good predictive performance and training time. Specifically, 32k is a training data size in which membrane potential errors from the test set are within the mean RMSE recording error (*∼* 4.8mV, averaged over pre-, mid-, post-activation periods) obtained from experimental neurons with multiple membrane potential recording data. For more details on generating simulated neuron training samples, see *Generating Training Data* under the *Methods* section.

To initially test EP-GAN performance, we evaluate EP-GAN predicted parameters for 200 simulated neurons outside of the training set (test set). To emulate *C. elegans*’ neuronal diversity, neurons in test set are divided into 3 response types - i) Transient outward rectifier type, ii) Outward rectifier type, and iii) Bistable type - that are currently found in non-spiking neurons of *C. elegans* according to their steady-state responses (Figure 2) [8, 16]. For a given neuron being evaluated, EP-GAN predicted HH-parameters are obtained as follows: For each training epoch, EP-GAN generates a set of HH-parameters for the neuron and at the end of the training, the parameter set which achieved the lowest Root Mean Square Error (RMSE) of membrane potential responses averaged across three time intervals - pre-activation (4, 5] seconds, mid-activation [5, 10] seconds, post-activation [10, 11) seconds - with respect to ground truths is reported (detailed descriptions of its calculation provided in supplementary and Figure 5A). In the case of multiple *N* -neurons being evaluated, the same procedure is followed except EP-GAN generates *N* -parameter sets in parallel at each epoch. Such multi-inference is possible due to EP-GAN being a neural network, where parallel processing of inputs can be done with minimal impact on inference speed. Using these procedures, EP-GAN predicted HH-parameters result in mean membrane potential RMSE error of **2.37mV** for test set (See Figure 2 for representative samples and Table S1 for the detailed breakdown of the errors). These errors are within the mean recording RMSE error of 4.8mV obtained from experimental neurons and their distributions were skewed uni-modal type where the majority of the errors fall within 4mV (Figure S1).

**Figure 2:**
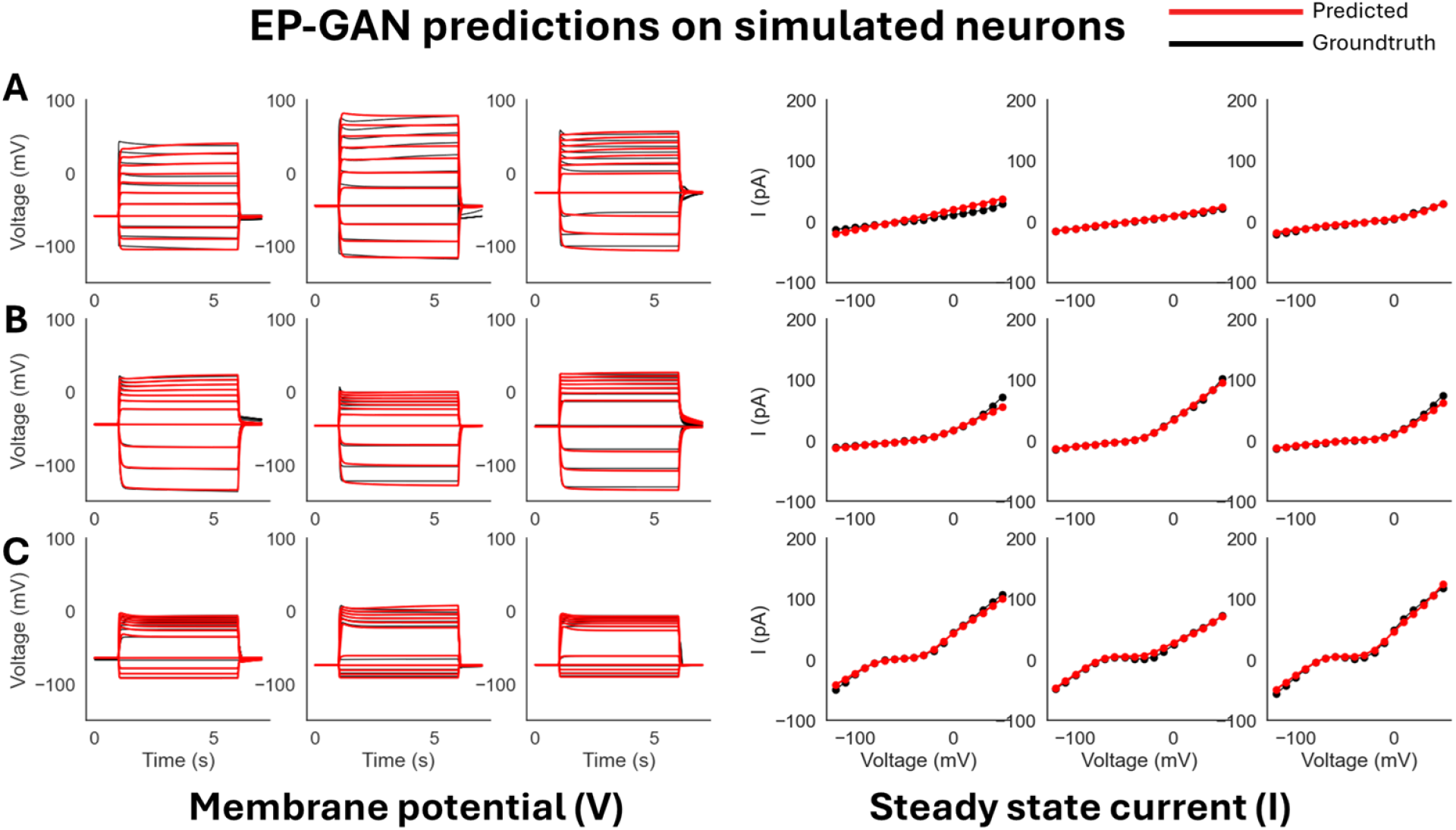
EP-GAN (32k) predictions on simulated neurons. **A**: EP-GAN predicted membrane potential traces and steady-state currents (Red) overlaid on top of groundtruth counterparts (black) for Transient outward rectifier neuron type. **B**: Outward rectifier neuron type. **C**: Bistable neuron type.

**Figure 5:**
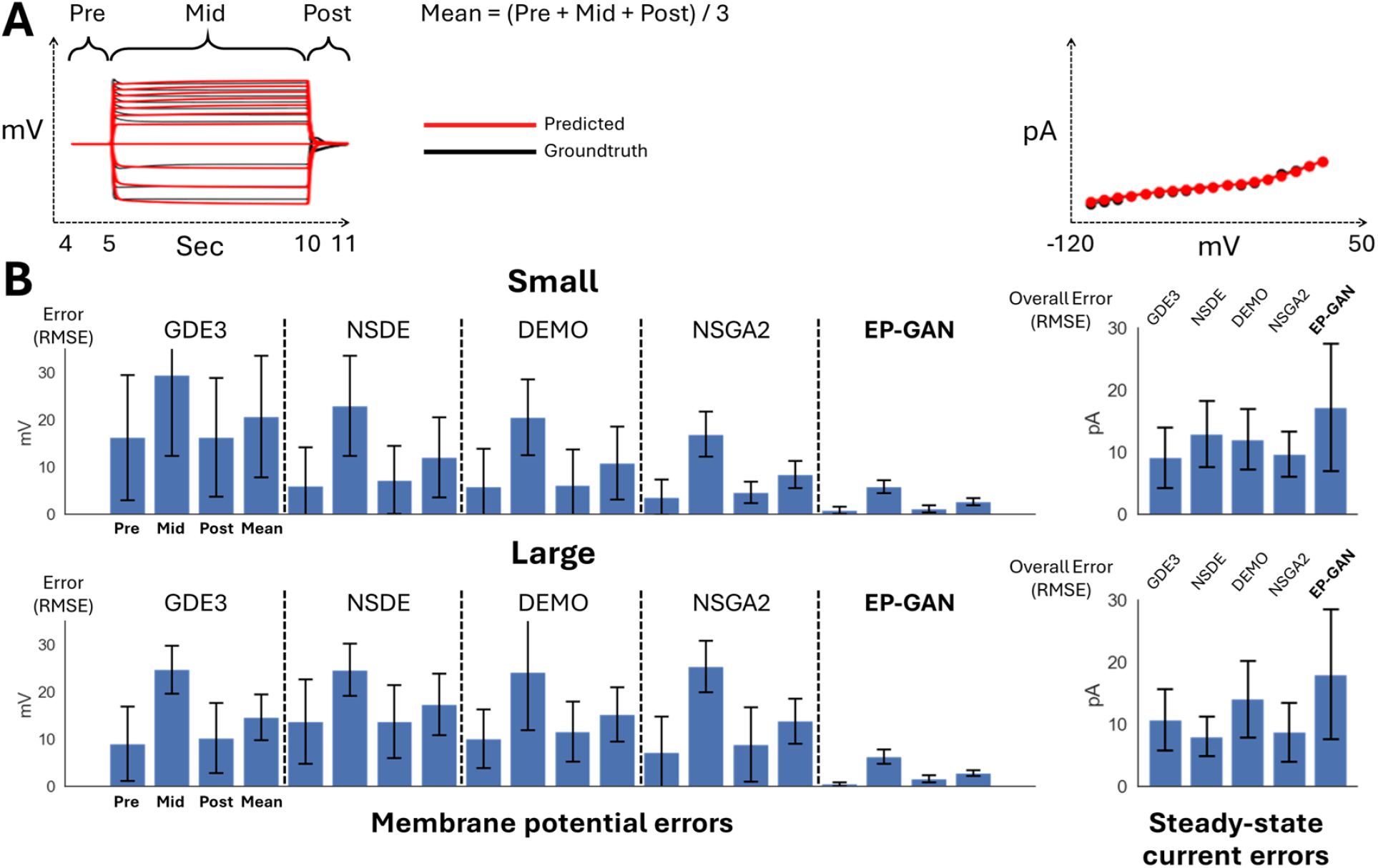
Bar plot showing the mean RMSE errors for membrane potential responses (pre-, mid-, post-activation periods, averaged error) and steady-state currents for 9 experimental neurons. **A**: Membrane potential responses (left) and steady-state currents (right) diagrams showing the time and voltage intervals of which the RMSE errors are computed. **B**: Bar plots showing RMSE errors for membrane potential responses and steady-state currents for small HH-model scenarios (Top) and Large HH-model prediction scenarios (Bottom). All methods use 32k sample size for both scenarios.

### Predictions on experimental neurons

We apply EP-GAN trained and tested on simulated data to predict HH-parameters for 9 experimentally recorded non-spiking neurons found in *C. elegans* - RIM, AFD, AIY, AWB, AWC, URX, RIS, DVC, HSN. Among these neurons, AWB, AWC, URX, RIS, DVC, HSN are novel recording data and were not previously modeled, whereas RIM, AIY and AFD neurons are publicly available and their modeling descriptions were elaborated by previous works [16, 22, 43]. Similar to prediction scenario on simulated neurons, we categorize experimental neurons into different response types according to their steady-state current responses. In particular, we classify (RIM, DVN, HSN) as transient outward rectifier type, (AIY, URX, RIS) as Outward rectifier type, and (AFD, AWB, AWC) as Bistable type.

We compare the performance of EP-GAN with 4 existing parameter inference methods: NSGA2, DEMO, GDE3, and NSDE. Unlike EP-GAN which can optimize multiple neurons in parallel, these are evolutionary methods where the optimization is done **from scratch** for each neuron. We therefore evaluate their respective performances relative to EP-GAN by normalizing on the *total number of simulated neuron samples* during the entire optimization task. Specifically, for all methods, we set the maximum number of neuron samples used during optimization to equal sizes. e.g., if EP-GAN is trained with 32k neuron samples to predict 9 neurons, NSGA2, DEMO, GDE3, and NSDE are each allocated with up to 3.5k samples during the search phase of HH-parameters for each neuron, thus adding up to a total of *∼* 32k samples for all 9 neurons. To test how the performance of each method scales with the amount of samples during optimization, we evaluate each method with both 32k and 64k total neuron samples.

The parameter selection process for NSGA2, DEMO, GDE3 and NSDE is as follows: During the search phase for each neuron, the parameter set candidates (i.e., population) are recorded at each iteration. At the end of the search phase, steady-state current profile of each parameter set candidate is evaluated with respect to the experimentally known bifurcation structure (i.e., *dI/dV*) of the neuron being inferred (e.g., bistable type). Upon evaluation, only the parameter sets satisfying the *dI/dV* bound constraints (*∼* 98% confidence interval) are kept. Such initial selection process is similar to the ones employed by previous methods utilizing evolutionary algorithms [16]. The *dI/dV* bound constraints are also used for generating EP-GAN training data (see *Generating Training Data* in *Methods* for more detail). The final parameter set is then chosen by selecting the one with the lowest RMSE membrane potential responses error averaged across pre-, mid- and post-activation periods identical to that of EP-GAN parameter selection process. For Multi-objective DE methods (DEMO, GDE3 and NSDE2), we follow the same configurations used in the literature to set their optimization scheme [16]. Specifically, we set crossover parameter CR and scale factor F to 0.3 and 1.5 respectively. For all 4 methods, NP (population size) is set to 600 with total 6 and 12 iterations (i.e., *∼*3.6*k* and *∼*6.2*k* samples per neuron for 32k and 64k total neuron samples respectively). For all methods, loss functions identical to the ones used for EP-GAN training (mean absolute errors, see *Methods* section for detail) are used for calculating the errors for membrane potential responses and steady-state currents for multi-objective optimization.

### Small HH-model scenarios (47 parameters)

We first test EP-GAN and existing methods with a “smaller” version of HH-model of 47 parameters where the individual channel parameters (*n* = 129) are frozen to default values given by [41]. The considered parameters consist of 16 conductance parameters of each channel (*g*_*Ch*_), 4 reversal potentials for calcium, potassium, sodium and leak channels (*V*_*Ca*_, *V*_*K*_, *V*_*Na*_, *V*_*L*_), 1 cell capacitance *C*, and 26 initial conditions for membrane potential *V*_0_ and channel activation variables (*m*_0_, *h*_0_). Such a parameter set is commonly targeted when fitting HH-models for individual neurons [7, 41]. For all methods, we test both 32k and 64k total sample sizes for the inference of 9 experimental neurons. Figure 3 illustrates that EP-GAN can reconstruct membrane potential responses close to ground truth responses. Indeed, upon inspecting the RMSE error for membrane potential responses (averaged over pre-, mid- and post-activation), EP-GAN (32k) median error of 2.5*mV* over 9 neurons is *∼* 50% lower than that of NSGA2 (64k) 4.3*mV* followed by DEMO (64k) 4.8*mV*, NSDE (64k) 5.5*mV* and GDE3 (64k) 6.0*mV* (Table 1, Table S2, Figure 5B). Among all 9 neurons being inferred, EP-GAN (32k) showed the best accuracy for HSN with 1.6*mV* and lowest accuracy for AFD with 4.9*mV*. With EP-GAN (64k), the median error further decreased (2.5*mV →* 2.4*mV*) where URX and RIS errors improved by 0.3*mV* and AFD error improved by 1.5*mV* over their 32k counterparts. Interestingly we note that the high accuracy of EP-GAN in predicting membrane potential is not necessarily complemented with steady-state currents (Table 1, Figure 5B). EP-GAN’s overall steady-state currents errors are generally higher than those of existing methods. A possible reason could be that the majority of these errors stems from lower and upper voltage ranges where the recording variations are high and thus could conflict with membrane potential responses optimization. Such competitive nature between membrane potential vs steady-state current optimizations has indeed been reported in previous works [16].

**Table 1:** Small HH-model scenarios RMSE errors for predicted membrane potential responses and steady-state currents. Each method is tested with 32k or 64k total sample sizes where the top row shows membrane potential responses RMSE errors averaged across pre-activation, mid-activation, post-activation periods and the bottom row shows steady-state currents RMSE errors across 18 voltage values.

| Method | Sample size | Median Error | RIM | DVC | HSN | AIY | URX | RIS | AFD | AWB | AWC |
| --- | --- | --- | --- | --- | --- | --- | --- | --- | --- | --- | --- |
| GDE3 | 32k | 16.9mV | 20.9 | 58.1 | 8.9 | 13.0 | 16.9 | 25.4 | 6.3 | 32.5 | 4.7 |
|  |  | 5.9pA | 5.8 | 5.8 | 19.8 | 4.8 | 2.4 | 5.9 | 19.1 | 7.2 | 11.9 |
|  | 64k | 6.0mV | 11.5 | 24.1 | 13.7 | 6.0 | 5.1 | 7.1 | 4.1 | 5.6 | 3.6 |
|  |  | 7.9pA | 7.5 | 4.6 | 6.8 | 7.2 | 18.7 | 7.9 | 42.1 | 17.6 | 46.7 |
| NSDE | 32k | 7.1mV | 38.7 | 8.3 | 20.2 | 5.7 | 7.1 | 11.0 | 5.5 | 6.3 | 6.6 |
|  |  | 13.6pA | 3.1 | 21.5 | 9.7 | 13.6 | 14.2 | 5.7 | 24.7 | 9.6 | 14.5 |
|  | 64k | 5.5mV | 15.4 | 8.7 | 20.2 | 13.5 | 5.2 | 5.5 | 4.9 | 4.9 | 4.0 |
|  |  | 9.7pA | 2.6 | 5.9 | 9.7 | 3.4 | 11.7 | 3.2 | 64.4 | 12.4 | 19.0 |
| DEMO | 32k | 6.7mV | 35.9 | 14.1 | 5.8 | 13.2 | 9.0 | 6.7 | 5.0 | 4.9 | 3.8 |
|  |  | 11.5pA | 6.5 | 13.8 | 14.6 | 5.2 | 5.4 | 9.7 | 23.8 | 11.5 | 18.7 |
|  | 64k | 4.8mV | 12.3 | 10.5 | 5.8 | 10.2 | 4.8 | 4.7 | **3.1** | 4.4 | 2.9 |
|  |  | 14.6pA | 4.4 | 6.5 | 14.6 | 4.1 | 15.2 | 10.3 | 41.0 | 23.1 | 17.8 |
| NSGA2 | 32k | 7.5mV | 12.4 | 15.4 | 2.6 | 9.8 | 6.1 | 6.4 | 7.5 | 9.4 | 6.6 |
|  |  | 10pA | 4.0 | 7.4 | 14.3 | 4.6 | 16.6 | 5.0 | 14.4 | 11.9 | 10.0 |
|  | 64k | 4.3mV | 10.5 | 19.0 | 5.2 | 13.3 | 3.5 | 3.2 | 4.2 | 4.3 | 3.1 |
|  |  | 8.4pA | 8.4 | 1.8 | 5.2 | 2.3 | 21.2 | 6.5 | 26.6 | 12.6 | 49.4 |
| **EPGAN**<br>**(Ours)** | 32k | 2.5mV | 3.4 | 2.4 | 1.6 | 2.5 | 3.2 | 1.7 | 4.9 | 2.5 | **2.0** |
|  |  | 13.8pA | 4.0 | 13.8 | 10.3 | 10.7 | 38.9 | 16.8 | 48.0 | 9.6 | 28.9 |
|  | 64k | **2.4mV** | **3.4** | **2.4** | **1.6** | **2.4** | **2.9** | **1.4** | 3.4 | **2.5** | 2.1 |
|  |  | 13.1pA | 3.2 | 13.9 | 2.5 | 9.8 | 16.5 | 13.1 | 49.7 | 11.6 | 27.9 |

**Figure 3:**
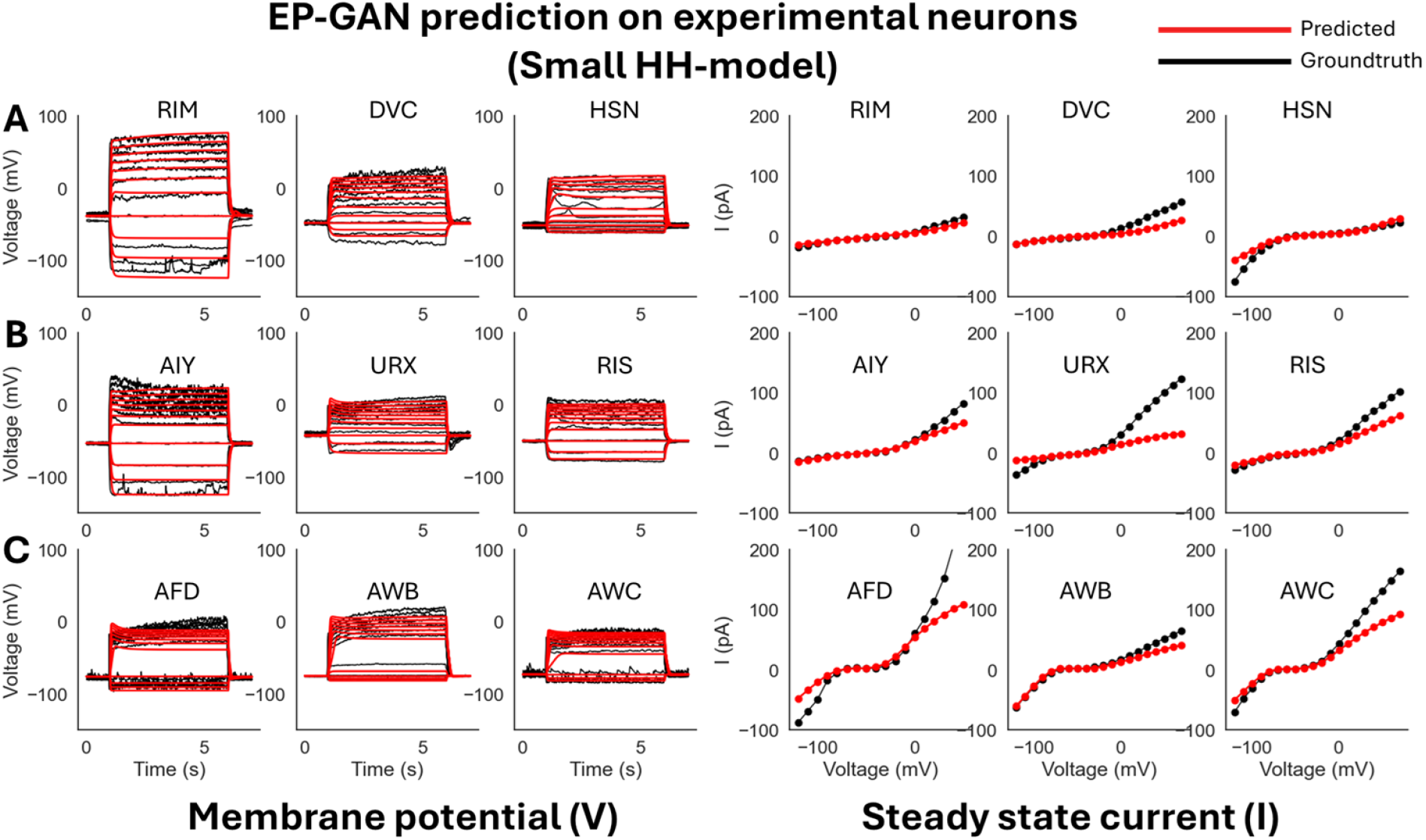
EP-GAN (32k) Prediction on experimental neurons (small HH-model). **A**: EP-GAN predicted membrane potential traces and steady-state currents (Red) overlaid on top of groundtruth counterparts (black) for Transient outward rectifier neuron type (RIM, DVC, HSN). **B**: Outward rectifier neuron type (AIY, URX, RIS). **C**: Bistable neuron type (AFD, AWB, AWC).

### Large HH-model scenarios (175 parameters)

We expand the domain of parameters being inferred by testing with respect to all 175 HH-model’s trainable parameters including the 47 parameters from the previous scenario + 129 channel parameters. The inclusion of channel parameters allows methods to further fine tune the HH-model. The minimum and maximum values for channel parameters are set to *±*50% from their default values (See *Generating Training Data* under *Methods* for more detail). Such a task introduces further challenges as the methods need to estimate *×*3 more parameters compared to the small HH-model scenario. From Figure 4, Figure 5B, Table 2, and Table S3 we see that while EP-GAN (32k) median membrane potential error increases slightly from 2.5*mV* to 2.7*mV*, its performance gaps over existing methods widen with *∼* 60% lower error than NSGA2 (64k) 7.5*mV*, followed by NSDE (64k) 8.6*mV* and DEMO (64k), GDE3 (64k) 10.5*mV*. EP-GAN trained for large HH-model also slightly improved overall steady-state current error (13.8*pA →* 12.4*pA*) alongside with membrane potential errors for RIM (3.4*mV →* 3.2*mV*) and URX (3.2*mV →* 3.0*mV*) indicating different selectivity for individual neurons for small vs large HH-model. Further increasing the sample size to 64k improved median errors of both membrane potential (2.7*mV →* 2.6*mV*) and steady-states responses (12.4*pA →* 8.6*pA*). Taken together, these results show EP-GAN predicting capabilities for HH-parameters with higher membrane potential responses accuracy and its ability to generalize to a larger parameter space with minimal performance loss.

**Table 2:** Large HH-model scenarios RMSE errors for predicted membrane potential responses and steady-state currents. Each method is tested with 32k or 64k total sample sizes where the top row shows membrane potential responses RMSE errors averaged across pre-activation, mid-activation, post-activation periods and the bottom row shows steady-state currents RMSE errors across 18 voltage values.

| Method | Sample size | Median Error | RIM | DVC | HSN | AIY | URX | RIS | AFD | AWB | AWC |
| --- | --- | --- | --- | --- | --- | --- | --- | --- | --- | --- | --- |
| GDE3 | 32k | 12.8mV | 14.0 | 12.5 | 12.8 | 19.4 | 9.0 | 15.2 | 29.4 | 10.8 | 9.2 |
|  |  | 9.6pA | 3.2 | 23.5 | 12.8 | 10.2 | 6.3 | 6.0 | 7.9 | 18.1 | 9.6 |
|  | 64k | 10.5mV | 14.0 | 11.0 | 10.5 | 11.7 | 12.4 | 9.0 | 5.0 | 9.5 | 4.7 |
|  |  | 4.9pA | 3.2 | 6.5 | 7.4 | 3.8 | 4.8 | 4.0 | 16.9 | 14.8 | 4.9 |
| NSDE | 32k | 16.1mV | 31.5 | 19.0 | 8.7 | 12.1 | 8.6 | 23.8 | 9.8 | 27.1 | 16.1 |
|  |  | 7.2pA | 7.8 | 8.0 | 9.6 | 6.8 | 5.1 | 4.2 | 18.2 | 7.2 | 6.2 |
|  | 64k | 8.6mV | 33.9 | 8.6 | 12.4 | 5.9 | 8.6 | 8.0 | 16.6 | 12.5 | 4.6 |
|  |  | 8.1pA | 4.0 | 29.6 | 8.1 | 4.7 | 5.1 | 4.3 | 9.1 | 15.0 | 14.9 |
| DEMO | 32k | 16.6mV | 28.0 | 17.8 | 16.6 | 10.2 | 22.9 | 18.1 | 6.0 | 13.1 | 5.1 |
|  |  | 11.9pA | 7.6 | 11.9 | 21.4 | 7.2 | 11.1 | 6.8 | 18.2 | 11.9 | 30.9 |
|  | 64k | 10.5mV | 10.5 | 29.5 | 11.0 | 10.2 | 6.8 | 18.4 | 6.0 | 13.1 | 4.4 |
|  |  | 8.0pA | 7.4 | 2.6 | 21.3 | 7.2 | 8.0 | 7.0 | 51.1 | 11.9 | 9.8 |
| NSGA2 | 32k | 13.4mV | 13.4 | 16.1 | 29.2 | 11.3 | 8.6 | 13.5 | 8.2 | 11.2 | 13.6 |
|  |  | 7.6pA | 6.6 | 7.6 | 5.8 | 8.7 | 5.1 | 2.7 | 24.1 | 10.9 | 7.6 |
|  | 64k | 7.5mV | 10.6 | 16.0 | 22.5 | 7.5 | 4.6 | 13.4 | 5.4 | 6.9 | 6.6 |
|  |  | 4.9pA | 4.9 | 3.2 | 3.7 | 1.2 | 9.5 | 3.1 | 24.7 | 13.7 | 6.9 |
| **EPGAN**<br>**(Ours)** | 32k | 2.7mV | 3.2 | **2.5** | 3.0 | 2.7 | **3.0** | 1.8 | 4.5 | 2.6 | **2.1** |
|  |  | 12.4pA | 3.2 | 12.4 | 17.4 | 10.5 | 36.6 | 21.9 | 43.2 | 9.3 | 8.4 |
|  | 64k | **2.6mV** | **3.2** | 2.9 | **2.5** | **2.6** | 3.4 | **1.8** | **4.4** | **2.6** | 2.2 |
|  |  | 8.6pA | 3.6 | 8.6 | 3.3 | 4.9 | 37.9 | 9.2 | 31.8 | 5.1 | 10.2 |

**Figure 4:**
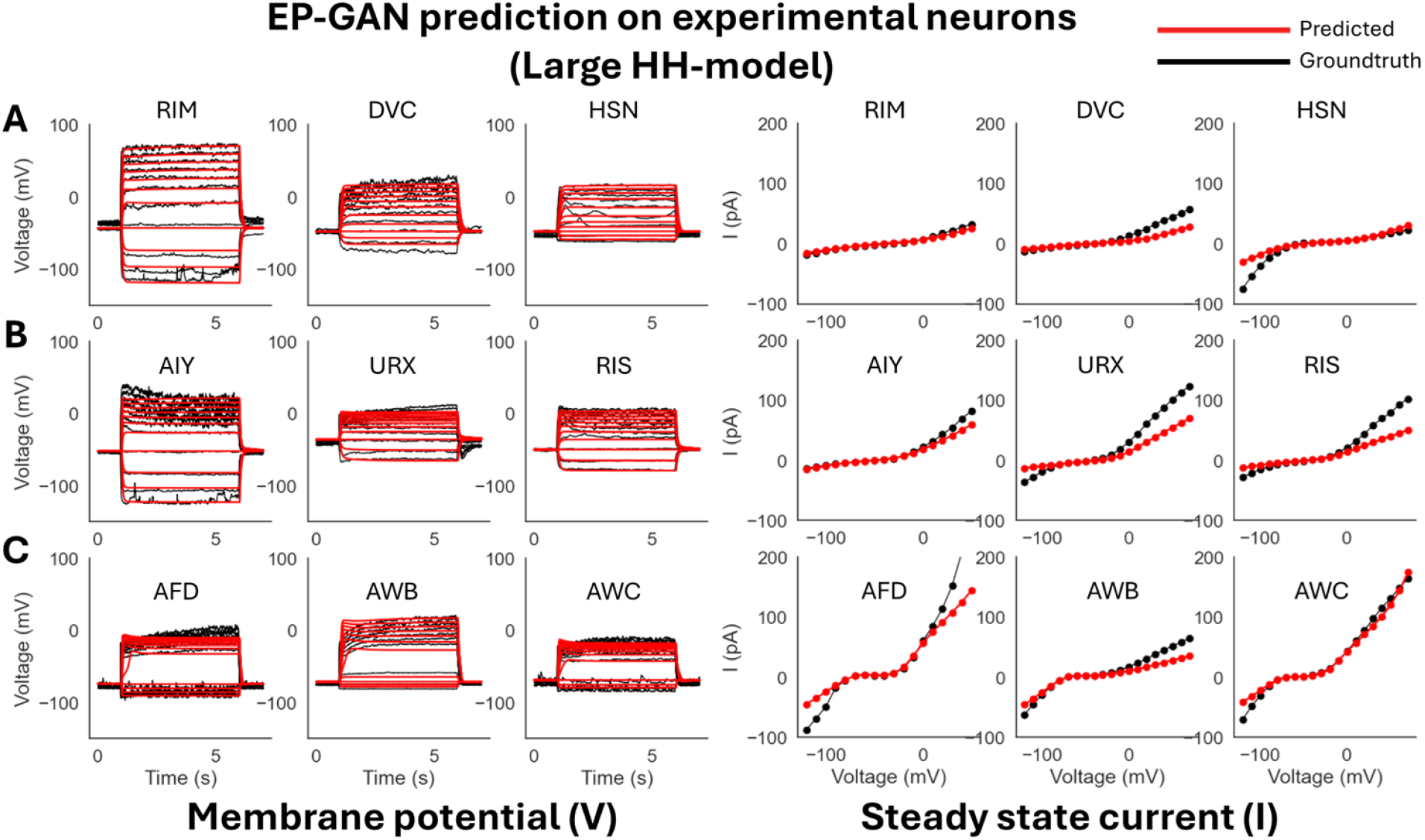
EP-GAN (32k) Prediction on experimental neurons (large HH-model). **A**: EP-GAN predicted membrane potential traces and steady-state currents (Red) overlaid on top of groundtruth counterparts (black) for Transient outward rectifier neuron type (RIM, DVC, HSN). **B**: Outward rectifier neuron type (AIY, URX, RIS). **C**: Bistable neuron type (AFD, AWB, AWC).

### Ablation studies

EP-GAN architecture also allows its membrane potential inputs to have arbitrary current-clamp protocol due to its RNN encoder component. To test the robustness of EP-GAN when incomplete input data is given, we provide the model with membrane potential responses and steady-state current inputs with missing data points. For each membrane potential responses and current profile, the data is reduced by 25%, 50%, and 75% each. For membrane potential responses data, the ablation is done on stimulus space where a 50% reduction corresponds to removing the upper half of the membrane potential responses traces each associated with a stimulus. For steady-state current profile, we remove the first *n*-data points where they are instead extrapolated using linear interpolation with existing data points.

Our results show that EP-GAN largely preserves accuracy even when both membrane potential and steady-state current inputs are masked (Figure 6, Table 3, Figure S4 (predicted steady-state currents)). In particular, EP-GAN preserves median membrane potential error (3.3*mV*) up to a 50% remaining in its inputs but becomes less accurate when up to 25% input remain (3.3*mV →* 5.4*mV*). Surprisingly, AFD neuron membrane potential error is improved when only 50% of input data is considered (5.2*mV →* 4.5*mV*). These results could be attributed to the random input masking employed during EP-GAN training (see *Methods* section for detail), which allows EP-GAN to make robust predictions even when conditioned with varying degrees of masked inputs.

**Table 3:**
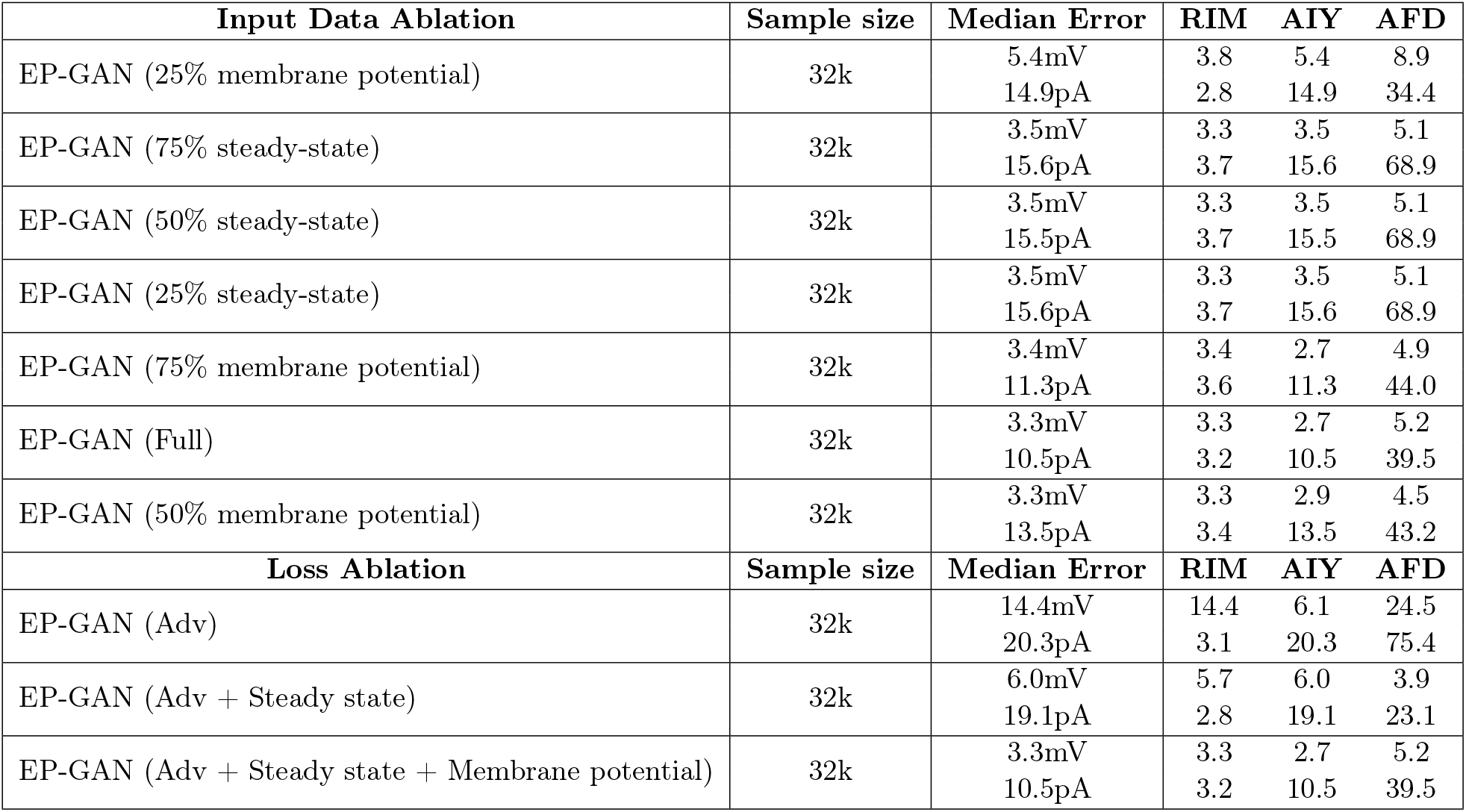
Ablation studies. **Top**: membrane potential responses and steady-state current errors achieved for EP-GAN (32k) when provided with incomplete input data. **Bottom**: membrane potential responses and steady-state current errors achieved for EP-GAN upon using only adversarial loss (Adv) and using adversarial + current reconstruction loss (Adv + Steady state) and all three loss components (Adv + Steady state + Membrane potential)

**Figure 6:**
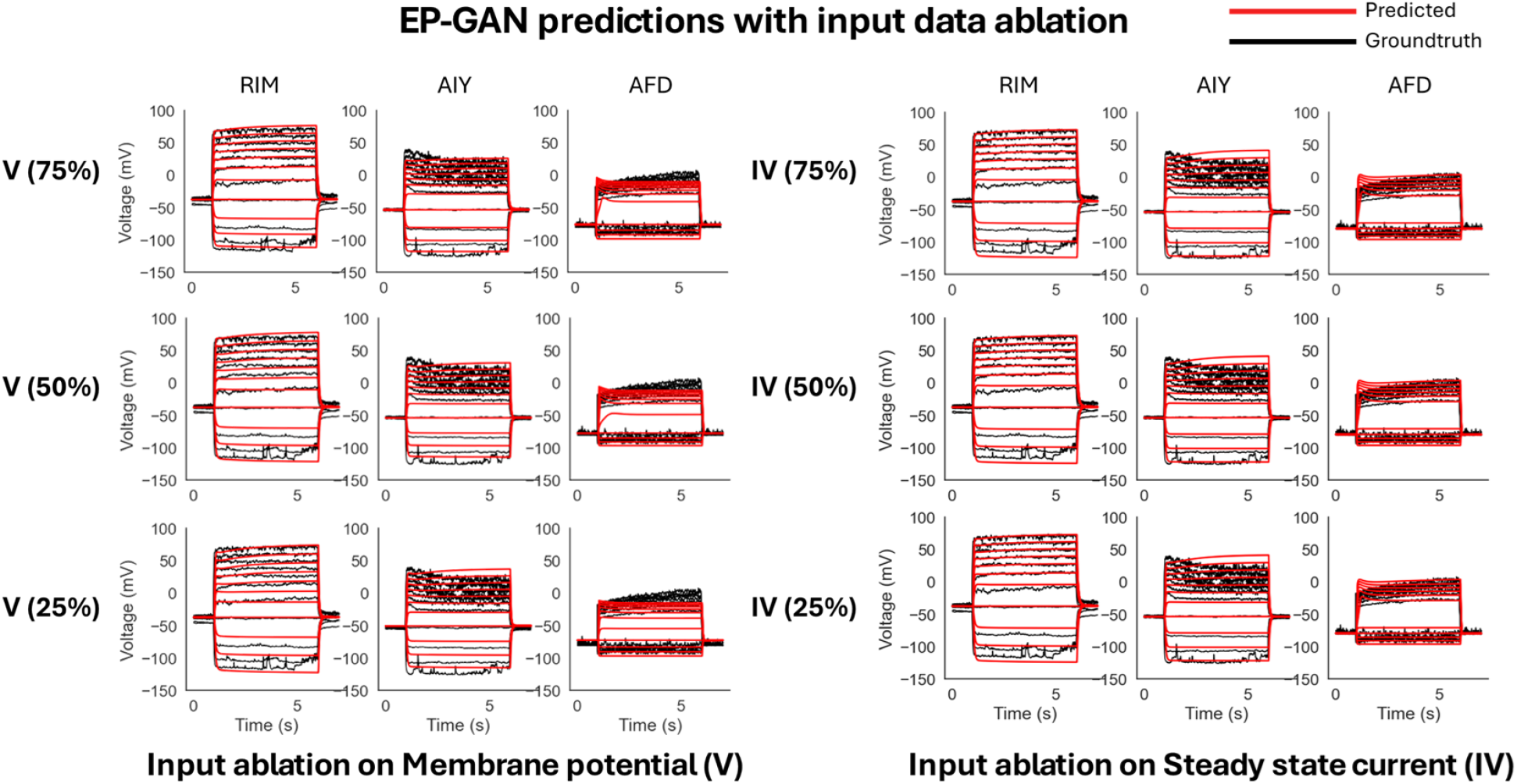
Input data ablation on EP-GAN (32k). **Left**: Reconstructed membrane potential responses for RIM, AIY and AFD when given with incomplete membrane potential responses data. Percentages in parenthesis represent the remaining portion of input membrane potential responses trajectories. **Right**: Reconstructed membrane potential responses for RIM, AIY, and AFD when given with incomplete steady-state current input.

We also performed ablation studies on model architecture by removing each loss component of the Generator module, allowing us to evaluate the relative contribution of each loss to accuracy. From Table 3 bottom, we see that removing the membrane potential loss term (V) results in a loss in performance for RIM and AIY but increase in accuracy for AFD. The result is consistent with input data ablation scenarios indicating AFD’s higher dependence on steady-state responses for good EP-GAN prediction. Upon removing the steady-state current reconstruction loss term (IV) in addition to the membrane potential reconstruction loss, we see further reduction in overall performance. These results highlight the significance of the reconstruction losses in aligning the Generator to produce the desired outputs.

### Parameter inference time

We also evaluate EP-GAN for its scalability by assessing its overall inference time and computational cost and comparing these to existing methods. Indeed, for estimation tasks involving many neurons, it is essential that the method is scalable so that the predictions are done within a reasonable time. In particular, for EP-GAN, the total time *T* needed for modeling *N* neurons including data generation and training time can be written as

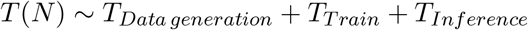

whereas for existing methods, the *T* (*N*) follows the form

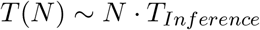

which increases linearly as *N* increases. Since *T*_*Inference*_ for EP-GAN is nearly instantaneous, it has a strong advantage in parameter prediction task involving multiple neurons. As an example, given a hypothetical task of modeling all 279 somatic neurons in the *C. elegans* nervous system, it would take DEMO, GDE3, NSDE or NSGA2 more than 44 days (assuming 7.2k samples per neuron, and our available computing environment) whereas, for EP-GAN, the process would be done within a day under a similar training setup. For a larger number of neurons, computational requirement of existing methods would grow linearly while EP-GAN would require constant time to complete the inference. Such scalability largely benefits from neural network being an inherently parallel architecture, allowing it to take multiple neuron profiles and output corresponding parameters in a single forward pass [44].

## Methods

We divide the methods section into two parts. In the first part we describe the detailed architecture of EP-GAN including its sub-modules with the simulation protocol used during training. In the second part, we describe the dataset and experimental protocol of novel neuron recording of AWB, AWC, URX, RIS, DVC, and HSN from *C. elegans* nervous system.

### Architecture of EP-GAN

#### Deep Generative Model for Parameter Prediction

EP-GAN receives neuronal recording data such as membrane potential responses and steady-state current profiles and generates a set of parameters that are associated with them in terms of simulating the inferred HH-model and comparing the simulated result with the inputs (Figure 1,7). We choose a deep generative model approach, specifically Generative Adversarial Network (GAN) as a base architecture of EP-GAN. The key advantage of GAN is in its ability to generate artificial data that closely resembles real data. The generative nature of GAN is advantageous for addressing the one-to-many nature of our problem, where there exists multiple parameter solutions for a given neuron recording. Indeed, several computational works attempting to solve inverse HH-model noted the ill-posed nature of the parameter solutions [5, 10, 11, 12, 13, 18, 45, 46]. Our approach is therefore leveraging GAN to learn a *domain of parameter sets* compatible with neuron recording instead of mapping directly onto a single solution. GAN consists of two separate networks, Generator and Discriminator. The goal of the Generator is to generate outputs that are indistinguishable from real data whereas the Discriminator’s goal is to distinguish outputs that are generated by the generator against real data. Throughout training, the Generator and the Discriminator engage in zero-sum game until they both converge to optimal states (i.e. Nash equilibrium) [47]. The particular architecture we use is Wasserstein GAN with gradient penalty (WGAN-GP), a variant of GAN architecture offering more stable training and faster convergence [48].

**Figure 7:**
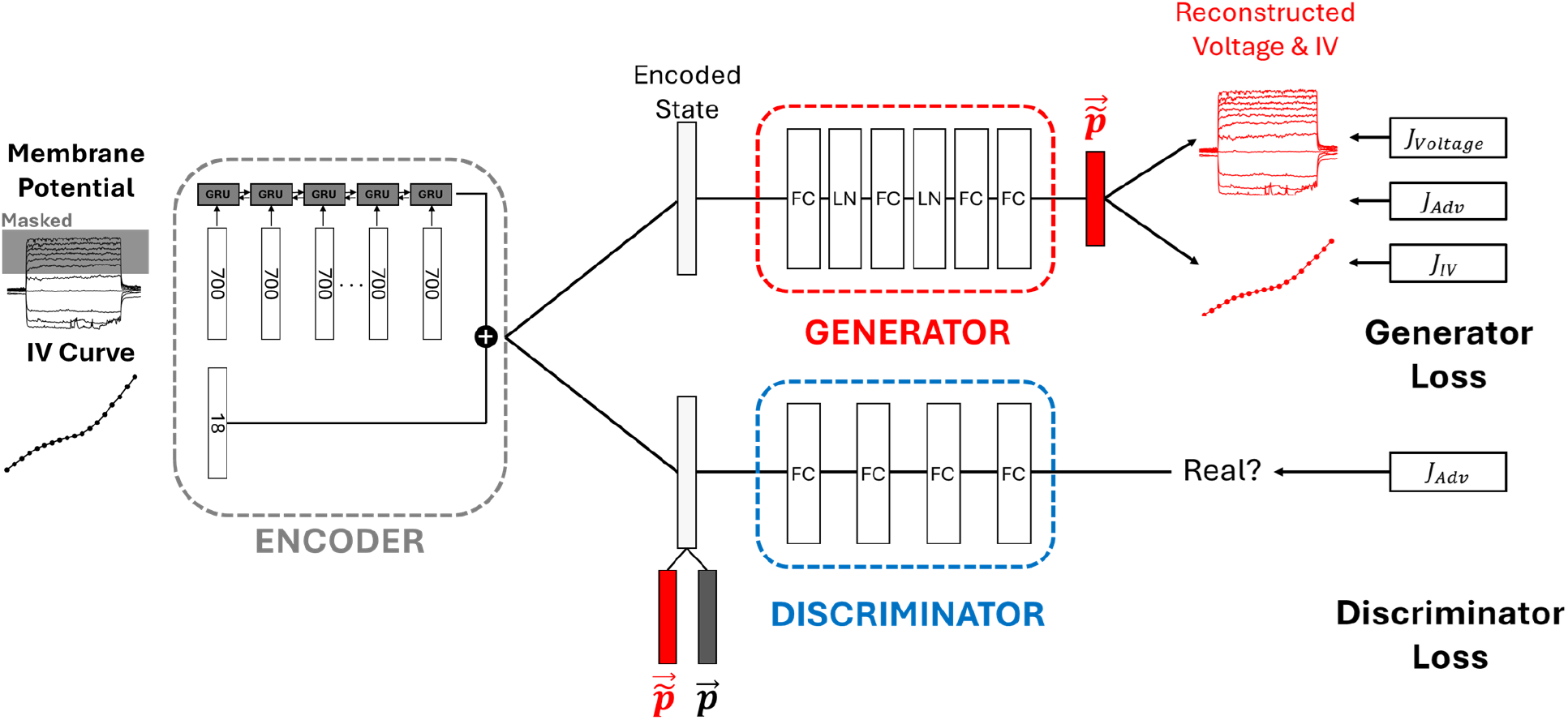
Architecture of EP-GAN. The architecture consists of an Encoder, Generator and Discriminator. Encoder compresses the membrane potential responses into a 1D vector (i.e., latent space) which is then concatenated with 1D steady-state current profile to be used as an input to both generator and discriminator. Generator translates the latent space vector into a vector of parameters 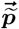 and the Discriminator outputs a scalar measuring the similarity between generated parameters 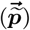 and ground truths 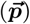. The Generator is trained with adversarial loss supplemented by reconstruction losses for both membrane potential responses and steady-state current profiles. The Discriminator is trained with discriminator adversarial loss only. Generator and Discriminator follow the architecture of Wasserstein GAN with gradient penalty (WGAN-GP) for more stable learning. During training, random masking is applied to input membrane potential responses which gradually decreases toward as the training continues.

#### Encoder Module

In addition to Generator and Discriminator, we implement Encoder module which pre-processes the input membrane potential responses for Generator and Discriminator. (Figure 7 left). Specifically, the encoder serves two roles: i) compression of membrane potential responses traces along the stimulus space, thus reducing its dimension from 2-dimensional to 1-dimensional, and ii) translation of membrane potential responses traces into a latent space encodinga meaningful internal representation for the Discriminator and Generator. The Encoder module uses Gated Recurrent Units (GRU) architecture, a variant of Recurrent Neural Network (RNN) to perform this task [49]. Each input sequence to a GRU cell at step *k* corresponds to the entire membrane potential response of size 350 (i.e., 350 time points, representing *t* = [4*s*, 11*s*]) concatenated with the associated stimulus trace of equal size of 350. Since GRU is agnostic to the number of steps in input sequence, such input structure allows EP-GAN to process a set of membrane potential traces with arbitrary current-clamp protocol. The output of the Encoder is a latent space vector of size 1024 encoding membrane potential responses information. The latent space vector is then concatenated with a 1D vector representing steady-state current profile which is used as an input to both Generator and Discriminator. During training, we randomly mask membrane potential traces to aid in better generalization in prediction [50, 51]. Specifically, we initially set the masking rate to 75% (i.e., 75% of membrane potential traces are randomly masked) and linearly decreases to 0% toward the end of training.

#### Discriminator Module

The goal of the Discriminator is given the input membrane potential responses and current profiles, to distinguish generated parameters from real ground truth parameters. The Discriminator receives as input the latent space vector from Encoder concatenated with generated or ground truth parameter vector and outputs a scalar representing the relative distance between two parameter sets (Figure 7, Eqn 1). Such a quantity is called Wasserstein distance or Wasserstein loss and differs from vanilla GAN Discriminator which only outputs 0 or 1. Wasserstein loss is known to remedy several common issues that arise from vanilla GAN such as vanishing gradient and mode collapse, leading to more stable training [48]. To further improve the training of WGAN architecture we supplement Wasserstein loss with a gradient penalty term, which ensures that the gradients of the Discriminator’s output with respect to the input has unit norms [52]. This condition is called Lipschitz continuity and prevents Discriminator outputs from having large variations when there is only small variations in the inputs [52, 53]. Combined together, the Discriminator is trained with the following loss:

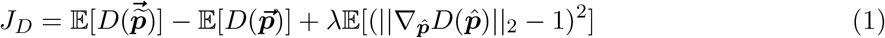

where 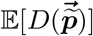 and 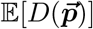 are the mean values of Discriminator outputs with respect to generated samples 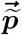 and real samples 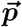 respectively and 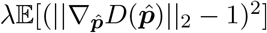 is the gradient penalty term modulated by *λ* where 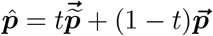 is the interpolation between generated and real samples with 0 *≤ t ≤* 1.

#### Generator Module

Being an adversary network of Discriminator, the goal of Generator is to fool the Discriminator by generating parameters that are indistinguishable from the real parameters. The Generator receives as input the concatenated vector from the Encoder and outputs a parameter vector (Figure 7). The module consists of 4 fully connected layers with layer normalization applied after the first two layers for improved model convergence [54]. Each parameter in the output vector is scaled between -1 and 1 which is then scaled back to the parameters’ original scales. The module is trained using 3 loss terms: i) Generator adversarial loss, ii) Membrane potential responses reconstruction loss and iii) Steady-state current reconstruction loss as follows:

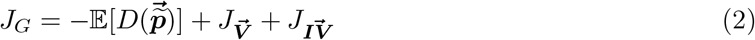

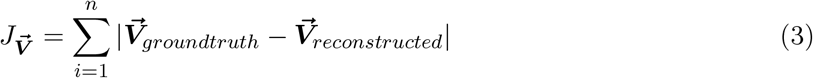

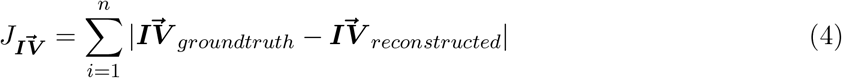

**Figure 8:**
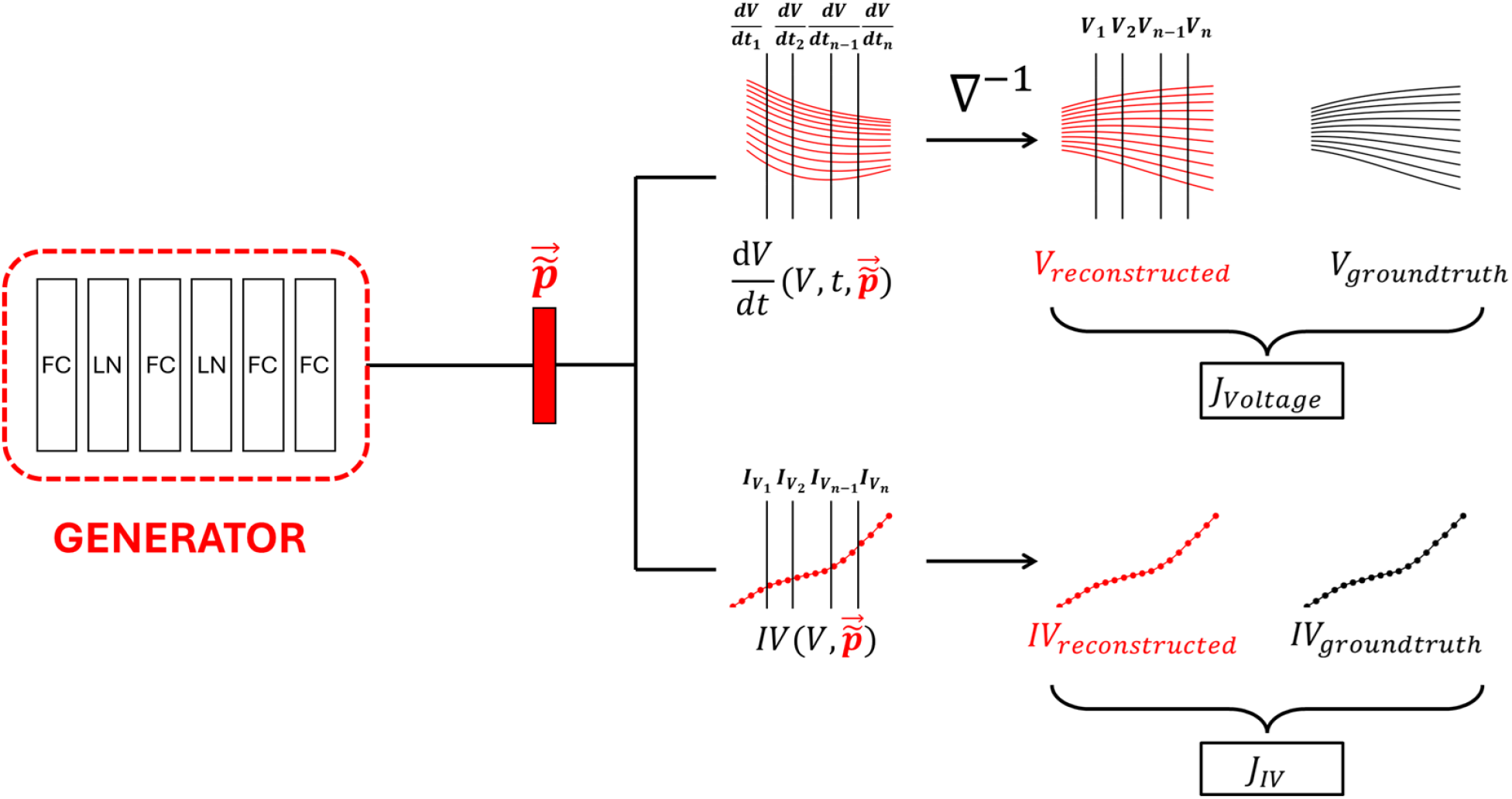
Description of membrane potential responses and current reconstruction losses for the Generator. Generated parameter vector 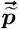 is used to evaluate membrane potential responses derivatives *dV/dt* at *n* time points sampled with fixed interval given the ground truth 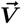 at those time points. The evaluated membrane potential responses derivatives are then used to reconstruct 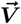 using the forward integration operation *∇*^−1^. The reconstructed 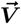 is then compared with ground truth 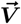 to evaluate the membrane potential responses reconstruction loss 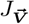. Current reconstruction is computed in a similar way via evaluating the currents at defined membrane potential responses steps *V* given generated parameters 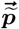 as inputs.

where 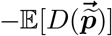 is Generator adversarial loss which is reciprocal of the mean Discriminator outputs with respect to generated samples and 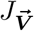 and 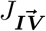 are *L*_1_ regression loss for reconstructed membrane potential responses and current profiles respectively. It’s important to note that 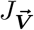 and 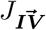 are part of Generator’s computation graph and thus force Generator to optimize them on top of adversarial loss (Figure 7). The composite loss function of Generator makes EP-GAN a “model-informed” GAN as HH-model itself becomes part of the training process. Such networks have shown to be more data efficient during training as they don’t rely solely on training data to learn effective strategy [55, 56]. The mathematical description of membrane potential responses and current reconstruction from generated parameter set 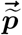 is as follows:

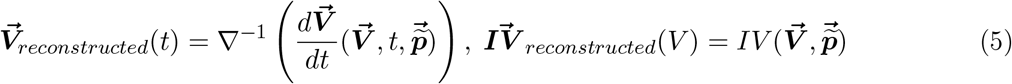

Here 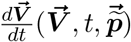 is the right-hand-side function of HH-model which computes the membrane potential responses derivative at time *t* given the membrane potential responses 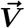 and parameter set 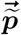 and 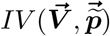 is the function that evaluates neuron’s current *I* given the membrane potential states 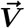 and parameter set 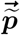. Membrane potential responses are reconstructed by first evaluating their derivatives with respect to ground truth membrane potential responses and generated parameters 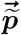 at regularly sampled time points. This is followed by forward integration operation *∇*^−1^ similar to Euler’s method to approximate 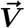 at sampled time points given the initial condition 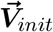:

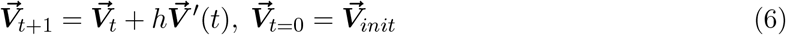

where *h* is the time interval between sampled derivatives. 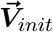 can be selected at any time point within the ground truth membrane potential state (e.g., pre-activation, mid-activation, post-activation) to reconstruct different membrane potential features. Notably, all computation steps consisting of forward integration process are expected to be differentiable. This is necessary to incorporate forward integration process as part of the generator network that requires full differentiability and thus trainable via back-propagation algorithm [57]. Computationally, we achieve this by manually implementing the forward integration process with discrete array operations that support auto-differentiation and vectorization (e.g. PyTorch Tensors) instead of simulating the membrane potential with ODE solvers [58]. The variable *h* can also be adjusted to increase the accuracy of the membrane potential responses reconstruction in exchange for the increased computational cost. The current profile is reconstructed by directly evaluating 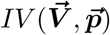 which uses the generated parameter set 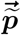 over the range of membrane potential responses values. We show in Table 3 that reconstruction losses are essential for the accuracy of predicted parameters.

#### Generating Training Data

For a successful training of a neural network model, the training data must be of sufficient number of samples, denoised, and diverse. To ensure these conditions are met with a simulated dataset, we employ a three-step process for generating training data (Figure 9).

**Figure 9:**
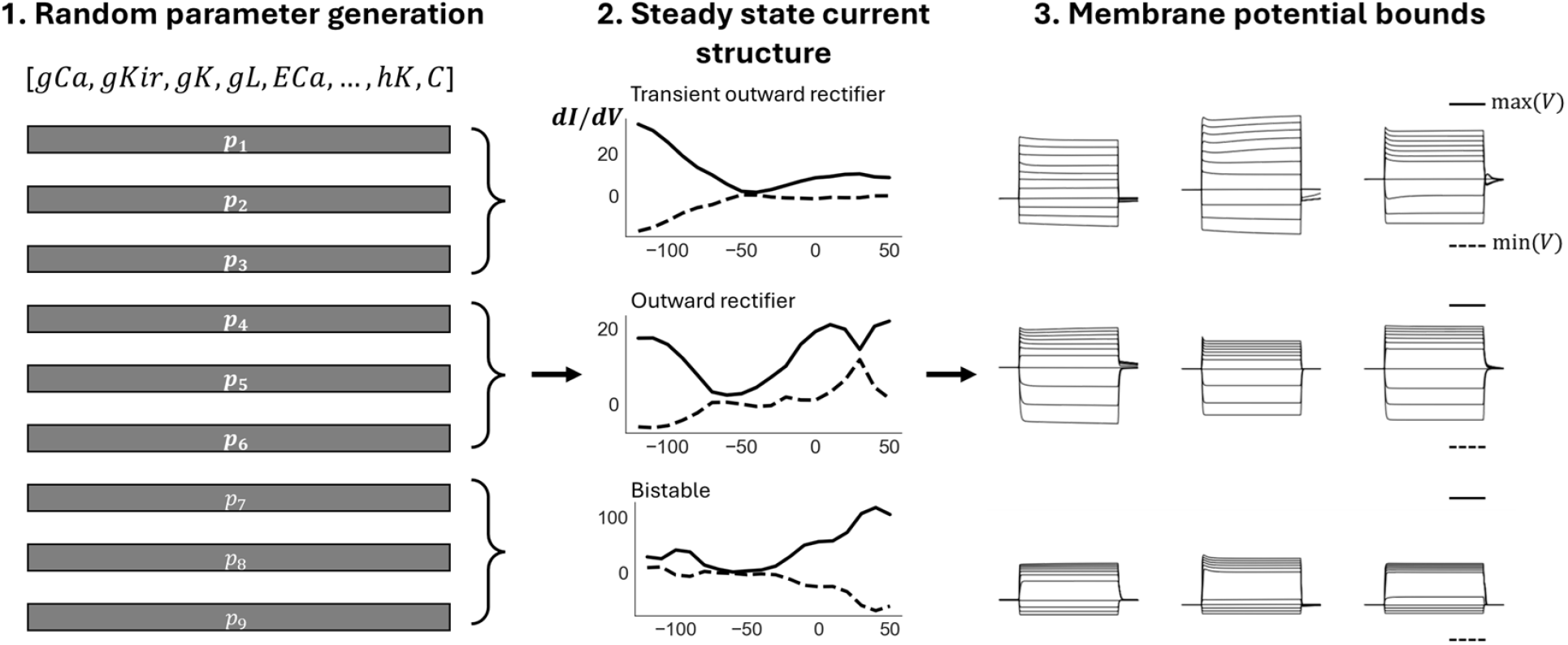
Training data generation. In Step 1, each parameter is initially sampled from biologically plausible ranges using both skewed gaussian (channel conductance) and uniform sampling. A parameter set consists of 175 parameters spanning 16 known ion channels in *C. elegans* and similar organisms. In Step 2 and 3, steady-state currents and membrane potential responses are evaluated for each set to ensure they satisfy the predefined constraints such as bifurcation structure represented by *dI/dV* bounds and minimum-maximum membrane potential across current-clamp protocols. Only parameter sets that meet both constraints are included in the training set.

**Step 1**. Randomly generate parameter sets by sampling each parameter within a pre-defined distribution. This distribution is skewed normal distribution for channel conductance parameters and uniform distributions for other parameters. The range is determined according to the biologically feasible ranges reported in the literature (See table *predicted parameters* in supplementary files for explicit range used for each parameter) [8, 22, 42]. In particular, search ranges for channel parameters are set to baseline *±*50% where baseline parameters are default parameters given by [41].

**Step 2**. Simulate steady-state current traces for each sampled parameter set followed by imposing bifurcation structure constraints on each of them. This is done by calculating the first derivative of the currents *dI/dV* with respect to voltage states and ensuring they are within the 98% confidence intervals of experimentally obtained *dI/dV* bounds. Specifically, we evaluate each parameter set with respect to the 3 neuron types that are found within *C. elegans* non-spiking neurons - Type 1: Transient outward rectifier (RIM, DVC, HSN), Type 2: Outward rectifier type (AIY, URX, RIS), and Type 3: Bistable type (AFD, AWB, AWC). During data generation, approximately same number of neurons are generated for each type to ensure balance between all neuron types. The step can be further extended with new neuron types by incorporating additional *dI/dV* bounds.

**Table 4:** Simulation protocols for simulated and experimental neurons.

| Neuron | Duration | Current-clamp<br>(min:step:max) | Stimulation period | Voltage-clamp<br>(min:step:max) |
| --- | --- | --- | --- | --- |
| Simulated | 15s | -15pA:5pA:35pA | 5s - 10s | -120mV:10mV:50mV |
| RIM | 15s | -15pA:5pA:35pA | 5s - 10s | -120mV:10mV:50mV |
| DVC | 15s | -2pA:1pA:8pA | 5s - 10s | -120mV:10mV:50mV |
| HSN | 15s | -2pA:1pA:8pA | 5s - 10s | -120mV:10mV:50mV |
| AIY | 15s | -15pA:5pA:35pA | 5s - 10s | -120mV:10mV:50mV |
| URX | 15s | -4pA:2pA:16pA | 5s - 10s | -120mV:10mV:50mV |
| RIS | 15s | -4pA:2pA:16pA | 5s - 10s | -120mV:10mV:50mV |
| AFD | 15s | -15pA:5pA:35pA | 5s - 10s | -120mV:10mV:50mV |
| AWB | 15s | -4pA:2pA:16pA | 5s - 10s | -120mV:10mV:50mV |
| AWC | 15s | -4pA:2pA:16pA | 5s - 10s | -120mV:10mV:50mV |

**Step 3**. Impose (minimum, maximum) constraints on the membrane potential response across current-clamp protocol. These values are set to (−100mV, 150mV) respectively for (−15pA, 35pA) current-clamp range. Parameter sets that do not satisfy the steady-state currents (Step 2) and membrane potential responses constraints are then removed from the training set. These constraints serve two purposes: i) remove parameter sets that result in non-realistic membrane potential responses/steady-state current profiles from the training set and ii) serve as an initial data augmentation process for the model. The constraints can also be extended or adjusted if deemed necessary for the improvement of the training process. Once constraints are applied, Gaussian noise is added to the membrane potential responses training data to mimic the measurement noises in experimental membrane potential responses recording data.

### Experimental Protocol

#### *C. elegans* Culture and Strains

All animals used in this study were maintained at room temperature (22-23°C) on nematode growth medium (NGM) plates seeded with E. coli OP50 bacteria as a food source [59]. Strains used in this study were: CX7893 kyIs405 (AWB), CX3695 kyIs140 (AWC), ZG611 iaIs19 (URX), EG1285 lin-15B(n765);oxIs12 (RIS), UL2650 leEx2650 (DVC) and CX4857 kyIs179 (HSN).

#### Electrophysiology

Electrophysiological recordings were performed on young adult hermaphrodites (*∼*3-days old) at room temperature as previously described [8]. The gluing and dissection were performed under an Olympus SZX16 stereomicroscope equipped with a 1X Plan Apochromat objective and widefield 10X eyepieces. Briefly, an adult animal was immobilized on a Sylgard-coated (Sylgard 184, Dow Corning) glass coverslip in a small drop of DPBS (D8537; Sigma) by applying a cyanoacrylate adhesive (Vetbond tissue adhesive; 3M) along one side of the body. A puncture in the cuticle away from the incision site was made to relieve hydrostatic pressure. A small longitudinal incision was then made using a diamond dissecting blade (Type M-DL 72029 L; EMS) along the glue line adjacent to the neuron of interest. The cuticle flap was folded back and glued to the coverslip with GLUture Topical Adhesive (Abbott Laboratories), exposing the neuron to be recorded. The coverslip with the dissected preparation was then placed into a custom-made open recording chamber (*∼*1.5 ml volume) and treated with 1 mg/ml collagenase (type IV; Sigma) for *∼*10 s by hand pipetting. The recording chamber was subsequently perfused with the standard extracellular solution using a custom-made gravity-feed perfusion system for *∼*10 ml.

All electrophysiological recordings were performed with the bath at room temperature under an upright microscope (Axio Examiner; Carl Zeiss, Inc) equipped with a 40X water immersion lens and 16X eyepieces. Neurons of interest were identified by fluorescent markers and their anatomical positions. Preparations were then switched to the differential interference contrast (DIC) setting for patch-clamp. Electrodes with resistance (RE) of 15-25 MΩ were made from borosilicate glass pipettes (BF100-58-10; Sutter Instruments) using a laser pipette puller (P-2000; Sutter Instruments) and fire-polished with a microforge (MF-830; Narishige). We used a motorized micromanipulator (PatchStar Micromanipulator; Scientifica) to control the electrodes back filled with standard intracellular solution. The standard pipette solution was (all concentrations in mM): [K-gluconate 115; KCl 15; KOH 10; MgCl2 5; CaCl2 0.1; Na2ATP 5; NaGTP 0.5; Na-cGMP 0.5; cAMP 0.5; BAPTA 1; Hepes 10; Sucrose 50], with pH adjusted with KOH to 7.2, osmolarity 320–330 mOsm. The standard extracellular solution was: [NaCl 140; NaOH 5; KCL 5; CaCl2 2; MgCl2 5; Sucrose 15; Hepes 15; Dextrose 25], with pH adjusted with NaOH to 7.3, osmolarity 330–340 mOsm. Liquid junction potentials were calculated and corrected before recording. Whole-cell current clamp and voltage-clamp experiments were conducted on an EPC-10 amplifier (EPC-10 USB; Heka) using PatchMaster software (Heka). Two-component capacitive compensation was optimized at rest, and series resistance was compensated to 50%. Analog data were filtered at 2 kHz and digitized at 10 kHz. Current-injection and voltage-clamp steps were applied through the recording electrode.

## Discussion

In this work, we introduce a novel deep generative method and system called ElectroPhysiomeGAN (EP-GAN), for estimating Hodgkin-Huxley model (HH-model) parameters given the recordings of neurons with graded potential (non-spiking). The proposed system encompasses RNN Encoder layer to process the neural recordings information such as membrane potential responses and steady-state current profiles and Generator layer to generate a large number of HH-model parameters in the order of 100s. The system can be trained entirely on simulation data informed by an arbitrary HH-model. When applied to neurons in *C. elegans*, EP-GAN generates parameters of HH-model which membrane potential responses are closer to ground truth responses than previous methods such as Differential Evolution and Genetic Algorithms. The advantage of EP-GAN is in the accuracy and inference speed achieved through fewer simulated samples than the previous methods for general parameter inference and is generic such that it doesn’t depend on the number of neurons for which inference is to be performed [7, 16, 41]. In addition, the method also largely preserves performance when provided with input data with partial information such as missing membrane potential responses (up to 50% missing) or steady-state current traces (up to 75% missing).

While EP-GAN is a step forward toward ElectroPhysiome model of *C. elegans*, its inability to support neurons with spiking membrane potential responses remains a limitation. The reason stems from the fact that neurons with spiking membrane potential responses are rare during the generation of training data of HH 16 ionic channels model without associated neuron channels, making their translation strategies to parameter space difficult to learn. A similar limitation is present with bi-stable membrane potential responses, e.g., AFD, AWB and AWC, although to a lesser extent. While the limitations for these profiles can be partially remedied through more training samples of their neuron types, their relative sparseness in the training data tends to cause lower quality of predicted parameters. Indeed, previous studies of *C. elegans* nervous system found that the majority of neurons exhibit graded membrane potential response instead of spiking [7, 21]. Furthermore, the limitation could also lie within the current architecture of EP-GAN as it currently processes data directly without a component that discerns and processes spiking membrane potential responses.

Improving the sampling strategy for training data alongside enhancement of network architecture could address these limitations in the future.

As discussed in the *Methods* section, it’s worth noting that EP-GAN does not necessarily recover the ground truth parameters that are associated with the input membrane potential responses and steady-state current profiles. This is mainly due to the fact that there may exist multiple parameter regimes for the HH-model which support the given inputs [5, 10, 11, 12, 13, 18, 45, 46]. The parameters generated by a single forward pass of EP-GAN (i.e., a single flow of information from the input to the output) could thus be interpreted as a one-time sampling from such a regime and a small perturbation to inputs may result in a different set of parameters. Such sensitivity to perturbation could be adjusted by supplementing the training samples or inputs with additional recording data (e.g., multiple recording data per neuron).

EP-GAN allows additional modifications to accommodate different configurations of the problem. For instance, update to HH-model would only require retraining of the network without changes to its architecture. Indeed, neuronal genome of *C. elegans* indicates additional voltage-gated channels that could be further incorporated to HH-models introduced in [7, 41] to improve its modeling accuracy of membrane potential dynamics [60]. Extending the inputs to include additional data, e.g., channel activation profiles, can also be done in a straightforward manner by concatenating them to the input vectors of the encoder network. Extending EP-GAN prediction capabilities to new neuron types can also be done by simply incorporating additional constraints during training data generation.

Despite its primary focus on *C. elegans* neurons, we believe EP-GAN and its future extension could be viable for modeling a variety of neurons in other organisms. Indeed, there are increasing advances in resolving connectomes of more complex organisms and techniques for large-scale neural activities [61, 62, 63, 64]. As neurons in these organisms can be described by a generic HH-model or similar differential equation model, EP-GAN is expected to be applicable and make contributions toward developing the biologically detailed nervous system models of neurons in these organisms.

## Acknowledgements

This work was supported in part by National Science Foundation grant CRCNS IIS-2113003 (JK,ES), Washington Research Fund (ES), CRCNS IIS-2113120 (QL), Kavli NSI Pilot Grant (QL), CityU New Research Initiatives/Infrastructure Support from Central APRC 9610587 (QL), the General Research Fund (GRF) and Early Career Scheme (ECS) Award from Hong Kong Research Grants Council RGC (CityU 21103522, CityU 11104123, CityU 11100524) (QL), and Chan Zuckerberg Initiative (to Cori Bargmann). Authors also acknowledge the partial support by the Departments of Electrical Computer Engineering (JK,ES), Applied Mathematics (ES), the Center of Computational Neuroscience (ES), and the eScience Institute (ES,JK) at the University of Washington. In addition, we thank Cori Bargmann and Ian Hope for *C. elegans* strains. Some strains were provided by the CGC, which is funded by NIH Office of Research Infrastructure Programs (P40 OD010440). We thank Saba Heravi for discussions regarding parameter inference for electrophysiological recordings.

## Notes

### Competing Interest Statement

The authors have declared no competing interest.

### Summary of Updates

Major revisions on Result and Methods section to address following aspects from the previous version: 1. Transient responses Made improvements to the training data generation and architecture of EP-GAN to improve its overall accuracy with predicted membrane potential responses. (Methods section, page 15) 2. RMSE time range choice for membrane potential baselines Supplemented the membrane potential regression loss with errors computed for 3 intervals (Figure 2, 3, Table S2, S3). 3. Introducing a richer set of metrics than RMSE Added ground truth vs predicted steady-state current responses plots into the main manuscript alongside with comparisons of membrane potential responses (Table 1,2,3, Figure 5, Table, S1, S2, S3) 4. Better comparison with existing methods Supplemented the optimization setup for existing methods (GDE3, NSDE, DEMO, and NSGA2) by incorporating steady-state response constraints as the initial selection process. Also expanded the testing scenarios by evaluating all methods w.r.t. both small and large HH-model estimation.

