## Supplementary Text for "ElectroPhysiomeGAN: Generation of Biophysical Neuron Model Parameters from Recorded Electrophysiological Responses"

### Supplementary Materials for ElectroPhysiomeGAN: Generation of Biophysical Neuron Model Parameters from Recorded Electrophysiological Responses

Jimin Kim[1], Minxian Peng[3], Shuqi Chen[3], Qiang Liu[3,*], and Eli Shlizerman[1,2,*]

[1]Department of Electrical and Computer Engineering, University of Washington
[2]Department of Applied Mathematics, University of Washington
[3]Department of Neuroscience, City University of Hong Kong
[*]Corresponding Authors

#### Error calculation

The overall errors between predicted membrane potential traces and their ground truth counterparts are computed using *root-mean-squared* error formula as follows

$$V_{error} = \sqrt{\frac{\sum_{t=1}^{T}\left(\sum_{k=1}^{N_{stim}}(V_{k,t}^{pred} - V_{k,t}^{gt})^2/N_{stim}\right)}{N_T}}$$

Voltage with upper subscripts *pred* and *gt* represents reconstructed membrane potential value at trace $k$ and time $t$ using predicted HH-parameters and ground truth variables respectively. $N_{stim}$ corresponds to a total number of stimulus value associated with a membrane potential trace. $N_T$ represents the total number of timepoints in which voltage traces are defined.

The error is computed for each of three intervals: pre-activation (4s - 5s], mid-activation [5s - 10s], and post-activation [10s - 11s), which are then averaged to compute the overall error.

#### Numerical simulation of HH-model

For efficient simulations of HH-model membrane potential dynamics, we use Julia ODE solver KenCarp47 algorithm supplemented by high-performance computing packages such as NumPy and SciPy [1, 2, 3]. Both relative and absolute tolerances for the ODE solver have been set to 1e-8 to ensure the accuracy of the simulations.

#### Recording data, Parameter units, ranges and predicted values

Electrophysiological recording data (current/voltage clamp) for 9 neurons used in experimental prediction scenarios can be found in supplementary files. Parameter ranges used for generating neuron samples for EP-GAN and existing methods, parameter trainability and their values predicted by EP-GAN can be found in *predicted parameters* table under supplementary files.

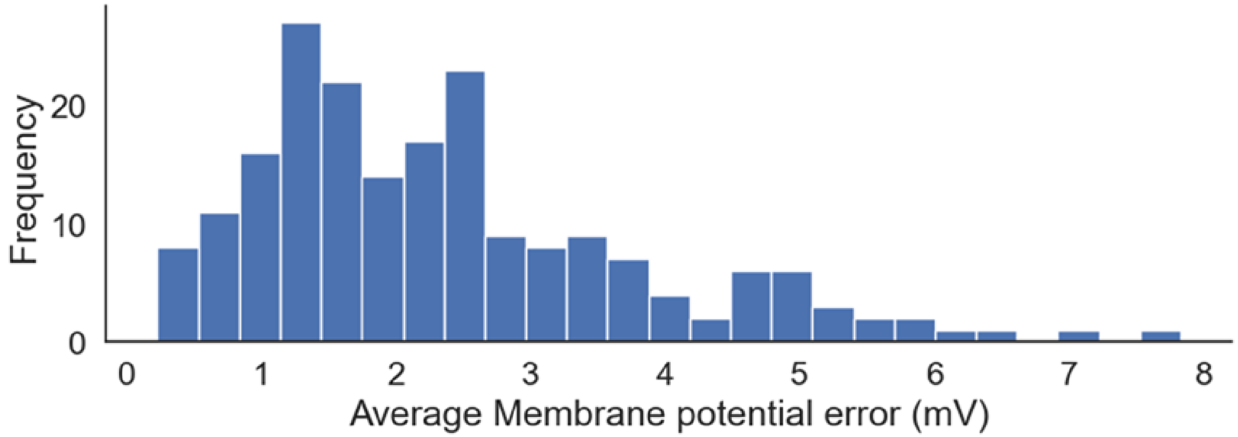

Figure. S1: RMSE Error distribution (averaged over pre-, mid-, post- activation time periods) for the simulated neurons ($n = 200$) in test set.

<table>
<tr><th>Method</th><th>Simulation #</th><th colspan="3">Test neurons</th></tr>
<tr><td rowspan="2">EPGAN</td><td rowspan="2">32k</td><td>1.33mV</td><td>3.63mV</td><td>2.15mV</td></tr>
<tr><td colspan="3">8.41pA</td></tr>
</table>

Table S 1: RMSE errors for membrane potential responses (top) and steady-state currents (bottom) for test neurons ($n = 200$) considered in *prediction on simulated neurons* scenario. Membrane potential responses errors are ordered as pre-activation error (4s - 5s), mid-activation error (5s - 10s), post-activation periods error (10s - 11s).

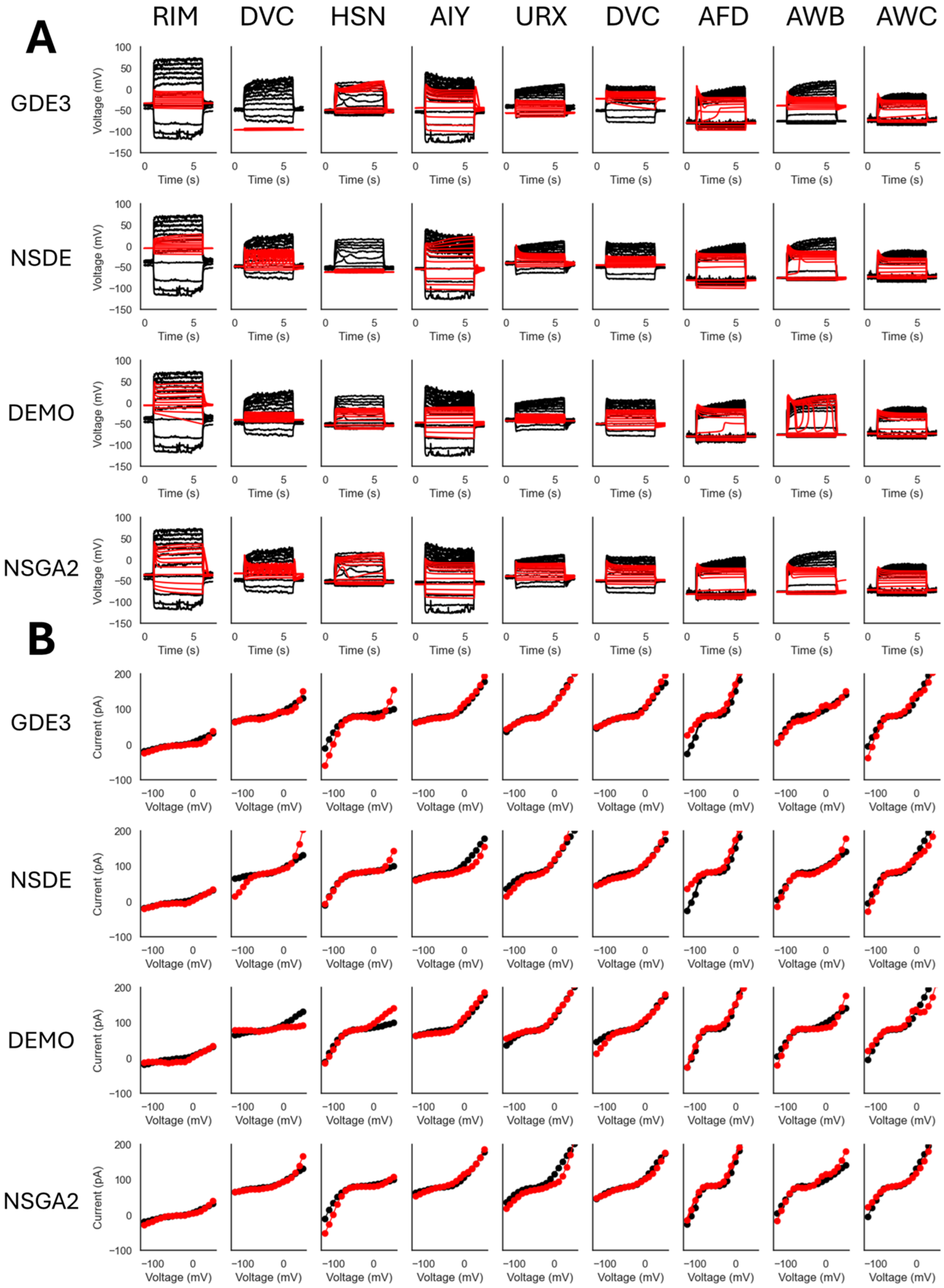

Figure. S2: **Small HH-model GDE3, NSDE, DEMO, NSGA2 Predictions (sample size = 32k) on experimental neurons**. **A**: Predicted membrane potential traces (Red) overlaid on top of ground truth (black) for all 9 experimental neurons. **B**: Predicted steady-state current traces (Red) overlaid on top of ground truth (black) for all 9 experimental neurons.

| Method | Neurons | | | | | | | | |
|---|---|---|---|---|---|---|---|---|---|
| GDE3 | **RIM** | | | **DVC** | | | **HSN** | | |
| | 5.63mV | 53.09mV | 3.92mV | 47.39mV | 79.42mV | 47.34mV | 2.8mV | 14.76mV | 8.98mV |
| | 5.78pA | | | 5.78pA | | | 19.84pA | | |
| | **AIY** | | | **URX** | | | **RIS** | | |
| | 9.75mV | 17.41mV | 11.75mV | 14.89mV | 25.19mV | 10.61mV | 28.39mV | 19.52mV | 28.18mV |
| | 4.82pA | | | 2.42pA | | | 5.94pA | | |
| | **AFD** | | | **AWB** | | | **AWC** | | |
| | 0.66mV | 17.37mV | 0.93mV | 36.72mV | 26.7mV | 34.21mV | 0.56mV | 12.42mV | 1.2mV |
| | 19.13pA | | | 7.22pA | | | 11.92pA | | |
| NSDE | **RIM** | | | **DVC** | | | **HSN** | | |
| | 33.29mV | 51.33mV | 31.54mV | 0.86mV | 19.88mV | 4.29mV | 9.76mV | 41.66mV | 9.22mV |
| | 3.14pA | | | 21.47pA | | | 9.7pA | | |
| | **AIY** | | | **URX** | | | **RIS** | | |
| | 0.46mV | 11.84mV | 4.69mV | 1.35mV | 15.19mV | 4.61mV | 5.86mV | 21.26mV | 5.96mV |
| | 13.63pA | | | 14.2pA | | | 5.68pA | | |
| | **AFD** | | | **AWB** | | | **AWC** | | |
| | 1.74mV | 13.16mV | 1.49mV | 0.5mV | 15.88mV | 2.39mV | 0.74mV | 17.47mV | 1.66mV |
| | 24.73pA | | | 9.55pA | | | 14.49pA | | |
| DEMO | **RIM** | | | **DVC** | | | **HSN** | | |
| | 32.73mV | 43.2mV | 31.84mV | 7.92mV | 27.06mV | 7.24mV | 1.18mV | 14.5mV | 1.59mV |
| | 6.52pA | | | 13.77pA | | | 14.63pA | | |
| | **AIY** | | | **URX** | | | **RIS** | | |
| | 7.26mV | 26.04mV | 6.27mV | 1.7mV | 22.14mV | 3.09mV | 1.29mV | 15.55mV | 3.26mV |
| | 5.16pA | | | 5.39pA | | | 9.68pA | | |
| | **AFD** | | | **AWB** | | | **AWC** | | |
| | 0.85mV | 13.06mV | 1.08mV | 0.05mV | 14.41mV | 0.21mV | 0.8mV | 9.44mV | 1.23mV |
| | 23.84pA | | | 11.47pA | | | 18.7pA | | |
| NSGA2 | **RIM** | | | **DVC** | | | **HSN** | | |
| | 3.1mV | 26.28mV | 7.88mV | 16.15mV | 21.63mV | 8.39mV | 0.48mV | 6.67mV | 0.64mV |
| | 3.97pA | | | 7.39pA | | | 14.29pA | | |
| | **AIY** | | | **URX** | | | **RIS** | | |
| | 3.58mV | 22.2mV | 3.71mV | 2.36mV | 10.37mV | 5.46mV | 2.88mV | 13.94mV | 2.36mV |
| | 4.59pA | | | 16.55 | | | 5.01 | | |
| | **AFD** | | | **AWB** | | | **AWC** | | |
| | 2.24mV | 17.99mV | 2.37mV | 0.52mV | 19.49mV | 8.32mV | 1.9mV | 14.82mV | 2.98mV |
| | 14.38pA | | | 11.86pA | | | 9.98pA | | |
| **EPGAN** | **RIM** | | | **DVC** | | | **HSN** | | |
| | 0.33mV | 8.23mV | 1.52mV | 0.19mV | 5.74mV | 1.22mV | 0.18mV | 4.02mV | 0.49mV |
| | 4.03pA | | | 13.8pA | | | 10.29pA | | |
| | **AIY** | | | **URX** | | | **RIS** | | |
| | 0.66mV | 6.41mV | 0.55mV | 1.66mV | 5.07mV | 2.82mV | 0.74mV | 3.77mV | 0.59mV |
| | 10.7pA | | | 16.84pA | | | 13.8pA | | |
| | **AFD** | | | **AWB** | | | **AWC** | | |
| | 3.15mV | 8.66mV | 2.87mV | 0.04mV | 7.23mV | 0.33mV | 0.66mV | 4.71mV | 0.72mV |
| | 47.97pA | | | 9.64pA | | | 28.86pA | | |

Table S 2: **Small HH-model RMSE errors (sample size = 32k) for membrane potential responses and steady-state currents for *predictions on small HH-model scenarios*.** For each neuron, top row shows the RMSE errors for 3 time intervals - pre-activation (4s - 5s), mid-activation (5s - 10s), and post-activation (10s - 11s) and bottom row shows the RMSE error for steady-state currents across 18 voltage points.

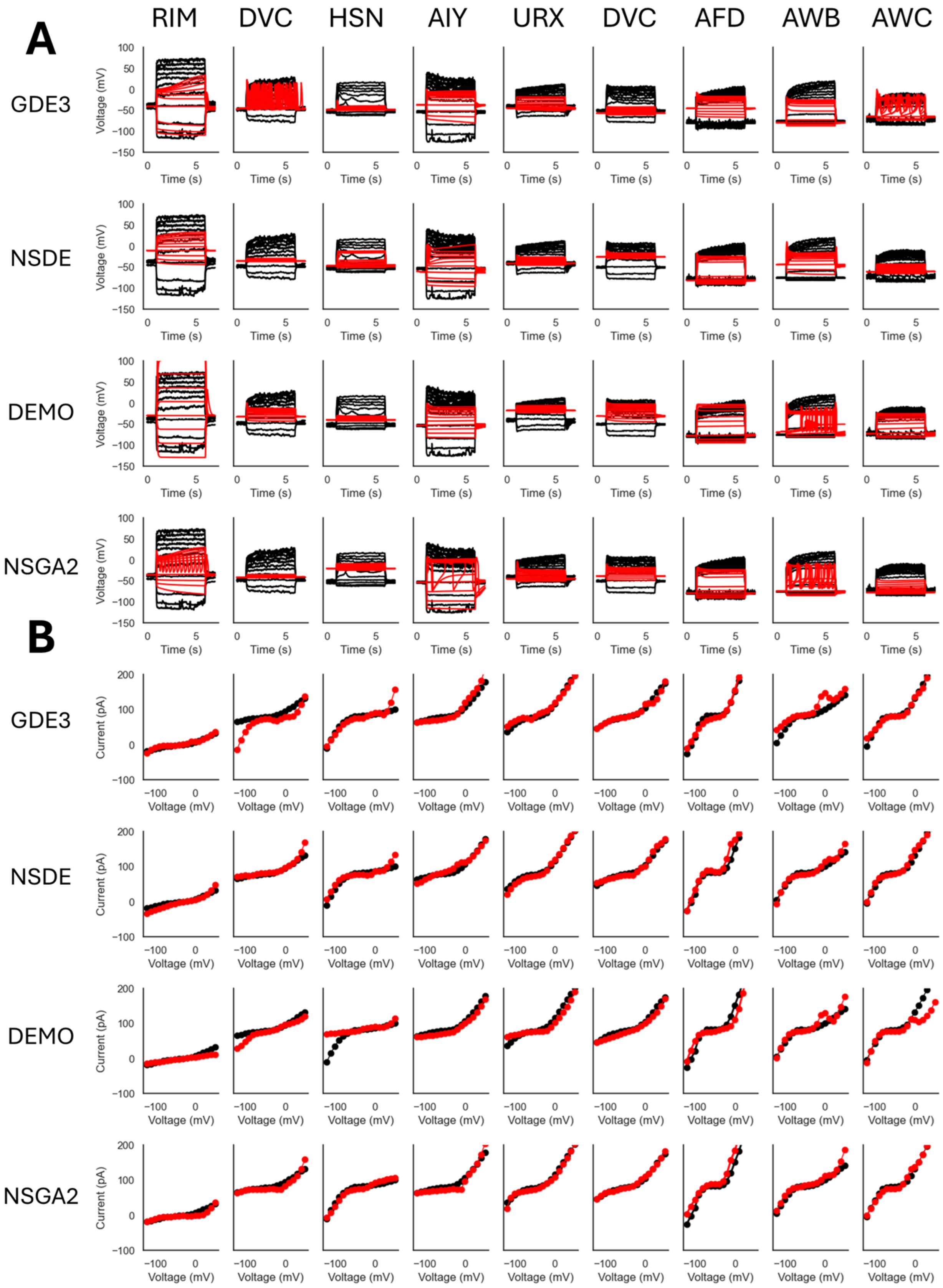

Figure. S3: **Large HH-model GDE3, NSDE, DEMO, NSGA2 Predictions (sample size = 32k) on experimental neurons**. **A**: Predicted membrane potential traces (Red) overlaid on top of ground truth (black) for all 9 experimental neurons. **B**: Predicted steady-state current traces (Red) overlaid on top of ground truth (black) for all 9 experimental neurons.

| Method | Neurons | | | | | | | | |
|---|---|---|---|---|---|---|---|---|---|
| | **RIM** | | | **DVC** | | | **HSN** | | |
| | 2.31mV | 30.01mV | 9.71mV | 3.08mV | 27.0mV | 7.54mV | 3.4mV | 31.0mV | 4.09mV |
| | 3.15pA | | | 23.53pA | | | 12.75pA | | |
| | **AIY** | | | **URX** | | | **RIS** | | |
| GDE3 | 16.88mV | 25.85mV | 15.51mV | 3.79mV | 20.27mV | 2.97mV | 6.54mV | 32.36mV | 6.61mV |
| | 10.24pA | | | 6.32pA | | | 5.99pA | | |
| | **AFD** | | | **AWB** | | | **AWC** | | |
| | 33.71mV | 20.5mV | 33.86mV | 3.8mV | 24.56mV | 4.14mV | 8.05mV | 11.58mV | 8.04mV |
| | 7.85pA | | | 18.08pA | | | 9.55pA | | |
| | **RIM** | | | **DVC** | | | **HSN** | | |
| | 27.97mV | 40.68mV | 25.98mV | 13.33mV | 30.91mV | 12.46mV | 5.03mV | 15.31mV | 5.73mV |
| | 7.75pA | | | 8.03pA | | | 9.58pA | | |
| | **AIY** | | | **URX** | | | **RIS** | | |
| NSDE | 6.81mV | 22.46mV | 7.05mV | 0.19mV | 21.14mV | 4.45mV | 24.54mV | 22.72mV | 24.22mV |
| | 6.82pA | | | 5.06pA | | | 4.17pA | | |
| | **AFD** | | | **AWB** | | | **AWC** | | |
| | 24.96mV | 2.09mV | 18.19mV | 31.41mV | 21.6mV | 28.39mV | 12.25mV | 22.77mV | 13.38mV |
| | 14.73pA | | | 7.24pA | | | 6.18pA | | |
| | **RIM** | | | **DVC** | | | **HSN** | | |
| | 8.91mV | 63.94mV | 11.11mV | 16.01mV | 21.32mV | 15.96mV | 11.8mV | 26.0mV | 12.11mV |
| | 7.57pA | | | 11.89pA | | | 21.4pA | | |
| | **AIY** | | | **URX** | | | **RIS** | | |
| DEMO | 1.46mV | 25.9mV | 3.21mV | 23.15mV | 18.16mV | 27.24mV | 19.32mV | 15.3mV | 19.8mV |
| | 7.24pA | | | 11.14pA | | | 6.81pA | | |
| | **AFD** | | | **AWB** | | | **AWC** | | |
| | 2.29mV | 11.92mV | 3.79mV | 7.0mV | 24.03mV | 8.3mV | 1.03mV | 10.97mV | 3.22mV |
| | 18.17pA | | | 11.9pA | | | 30.85pA | | |
| | **RIM** | | | **DVC** | | | **HSN** | | |
| | 3.58mV | 32.89mV | 3.63mV | 7.55mV | 33.6mV | 7.06mV | 31.5mV | 24.36mV | 31.81mV |
| | 6.57pA | | | 7.63pA | | | 5.75pA | | |
| | **AIY** | | | **URX** | | | **RIS** | | |
| NSGA2 | 0.38mV | 14.99mV | 18.39mV | 3.61mV | 20.4mV | 1.87mV | 11.87mV | 17.09mV | 11.45mV |
| | 8.68pA | | | 5.13pA | | | 2.74pA | | |
| | **AFD** | | | **AWB** | | | **AWC** | | |
| | 0.8mV | 22.68mV | 1.14mV | 1.14mV | 30.38mV | 2.11mV | 5.12mV | 32.47mV | 3.17mV |
| | 24.07pA | | | 10.9pA | | | 7.55pA | | |
| | **RIM** | | | **DVC** | | | **HSN** | | |
| | 0.24mV | 7.78mV | 1.65mV | 0.30mV | 5.90mV | 1.39mV | 0.46mV | 6.62mV | 2.02mV |
| | 3.21pA | | | 12.35pA | | | 17.44pA | | |
| | **AIY** | | | **URX** | | | **RIS** | | |
| **EPGAN** | 0.43mV | 6.57mV | 1.12mV | 0.74mV | 4.58mV | 3.78mV | 0.27mV | 3.42mV | 1.78mV |
| | 10.52pA | | | 36.64pA | | | 21.86pA | | |
| | **AFD** | | | **AWB** | | | **AWC** | | |
| | 1.68mV | 9.99mV | 1.86mV | 0.37mV | 7.24mV | 0.31mV | 0.57mV | 4.93mV | 0.84mV |
| | 43.20pA | | | 9.27pA | | | 8.35pA | | |

Table S 3: **Large HH-model RMSE errors (sample size = 32k) for membrane potential responses and steady-state currents for *predictions on large HH-model scenarios*.** For each neuron, top row shows the RMSE errors for 3 time intervals - pre-activation (4s - 5s), mid-activation (5s - 10s), and post-activation (10s - 11s) and bottom row shows the RMSE error for steady-state currents across 18 voltage points.

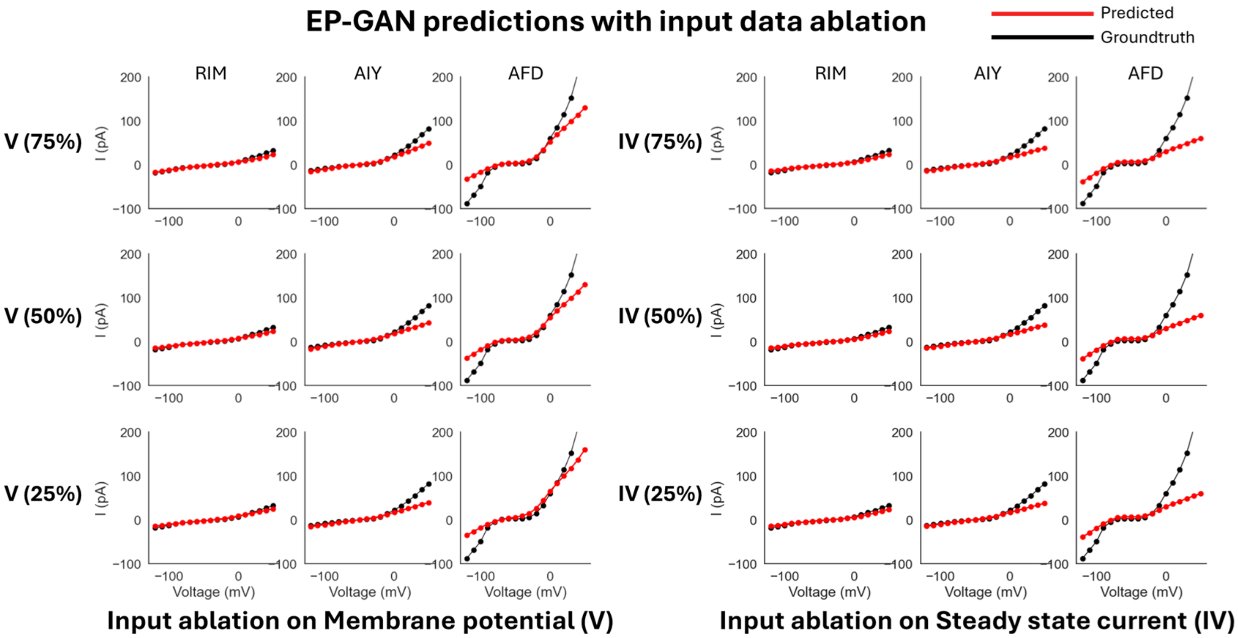

Figure. S4: **Input data ablation on EP-GAN (32k)**. **Left**: Reconstructed steady-state currents when given with incomplete membrane potential responses data. Percentages in parenthesis represent the remaining portion of input membrane potential responses trajectories. **Right**: Reconstructed steady-state currents when given with incomplete steady-state current input.
